# Coherent theta activity in the medial and orbital frontal cortices encodes reward value

**DOI:** 10.1101/2020.09.22.308809

**Authors:** Linda M. Amarante, Mark Laubach

**Affiliations:** Department of Neuroscience, American University, Washington, DC, USA

## Abstract

This study examined how the medial frontal (MFC) and orbital frontal (OFC) cortices process reward information. We simultaneously recorded local field potentials in the two areas as rats consumed liquid sucrose rewards. Both areas exhibited a 4-8 Hz “theta” rhythm that was phase locked to the lick cycle. The rhythm tracked shifts in sucrose concentrations and fluid volumes, demonstrating that it is sensitive to differences in reward magnitude. The coupling between the rhythm and licking was stronger in MFC than OFC and varied with response vigor and absolute reward value in the MFC. Spectral analysis revealed zero-lag coherence between the cortical areas, and found evidence for a directionality of the rhythm, with MFC leading OFC. Our findings suggest that consummatory behavior generates simultaneous theta range activity in the MFC and OFC that encodes the value of consumed fluids, with the MFC having a top-down role in the control of consumption.

## INTRODUCTION

The medial and orbital frontal cortices (MFC and OFC) are two of the most studied parts of the cerebral cortex for their role in value-guided decision making, a process that ultimately results in animals consuming rewarding foods or fluids. There are extensive anatomical connections between the various parts of the MFC and OFC in rodents (Gabbott et al., 2003; Gabbott et al., 2005; Barreiros et al., 2020), and the regions are part of the medial frontal network (Öngür and Price, 2000). The MFC and OFC are thought to have specific roles in the control of behavior and specific homologies with medial and orbital regions of the primate frontal cortex (MFC: Laubach et al., 2018; OFC: Izquierdo, 2017). The extensive interconnections between MFC and OFC suggest that the two regions work together to control value-guided decisions. Unfortunately, few, if any, studies have examined concurrent neural processing in these regions of the rodent brain as animals perform behavioral tasks that depend on the two cortical regions.

In standard laboratory tasks, the action selection and outcome evaluation phases of value-guided decisions are commonly conceived as separate processes (Rangel et al., 2008). MFC and OFC may contribute independently to these processes or interact concurrently across them. Though there is some variation across published studies, most argue for MFC having a role in action-outcome processing (Alexander and Brown, 2011; Simon et al., 2015) and OFC having a role in stimulus-outcome (stimulus-reward) processing (Gallagher et al., 1999; Schoenbaum and Roesch, 2005; Simon et al., 2015). The present study directly compared neural activity in the MFC and OFC of rats as they performed a simple consummatory task, called the Shifting Values Licking Task, or SVLT (Parent et al., 2015a). Importantly, the task depends on the ability of animals to guide their consummatory behavior based on the value of available rewards, and performance of these kinds of tasks depends on both the MFC (Parent et al., 2015a,b) and OFC (Kesner and Glibert, 2007). The goal of the study was to use the SVLT to determine if the MFC and OFC have distinct roles in processing reward information, e.g. varying with action (licking) in MFC and the sensory properties of the rewards in OFC.

Most published studies on reward processing used operant designs with distinct actions preceding different outcomes. For example, a rat might respond in one of two choice ports to produce a highly valued reward, delivered from a separate reward port. To collect the reward, the rat has to travel across an operant chamber and then collect a food pellet or initiate licking on a spout to collect the reward. In such tasks (Pratt and Mizumori, 2001; van Durren et al., 2009; van Wiingerden et al., 2010; Riceberg and Shapiro, 2017; Jarovi et al., 2018; Siniscalchi et al., 2019), neural activity during the period of consumption might reflect the properties of the reward, how the animal consumes it, and/or the behaviors that precede reward collection (e.g. locomotion). As such, it is difficult to isolate reward specific activity using such operant designs.

Several published studies have used simpler consummatory and Pavlovian designs, and found neural activity in the MFC is selectively modulated during active consumption (Petykó et al., 2009; Horst and Laubach, 2013; Petykó et al., 2015). None of these tasks used fluids with different reward values. Amarante et al. (2017) was the first study to examine if similar neural activity was associated with animals consuming different magnitudes of reward. The study used the SVLT and presented rats with rewards that differed in terms of the concentration of sucrose contained in the rewarding fluids. The study found that neural activity in the MFC is entrained to the animals’ lick cycle and the strength of entrainment varies with the value of the rewarding fluid, i.e. stronger entrainment with higher value reward. The study also used reversible inactivation methods to demonstrate that licking entrainment depends on the MFC.

In the present study, we used the SVLT, and several variations on the basic task design, to study consumption related activity in MFC and OFC. Spectral analyses were used to account for the extent to which neural activity in each area was entrained to licking and if there was evidence for directionality of lick-entrainment among sites in the MFC and OFC. A custom designed syringe pump was used to deliver different volumes of fluid over a common time period (Amarante et al., 2019). This device allowed us to directly compare neural activity associated with differences in sucrose concentration and fluid volume. We further manipulated the predictability of changes in reward magnitude to assess how predictable and unpredictable rewards are processed and used a third, intermediate level of reward to assess if reward magnitudes are encoded in a relative or absolute manner. Our findings suggest that both areas encode the value of consumed fluids and that the MFC may have a top-down role in coordinating reward processing.

## RESULTS

### Shifting Values Licking Task: Effects of reward magnitude on consummatory behavior

The Shifting Values Licking Task (Amarante et al., 2017; Figure 1A) was used to assess reward encoding across the MFC and OFC as 12 rats experienced shifts in reward value defined by differences in sucrose concentration or fluid volume. Shifts in concentration were between 16% and 4% sucrose in a volume of 30 μL. Shifts in volume were between 30 μL and 10 μL containing 16% sucrose. Concentrations and volumes alternated over periods of 30 sec (Figure 1B, left). LFP activity was recorded from 16-channel multi-electrode arrays in the MFC in 10 of the 12 rats and OFC in 6 of the 12 rats (recording locations are shown in Figure 1 - figure supplement 1).

**Figure 1.**
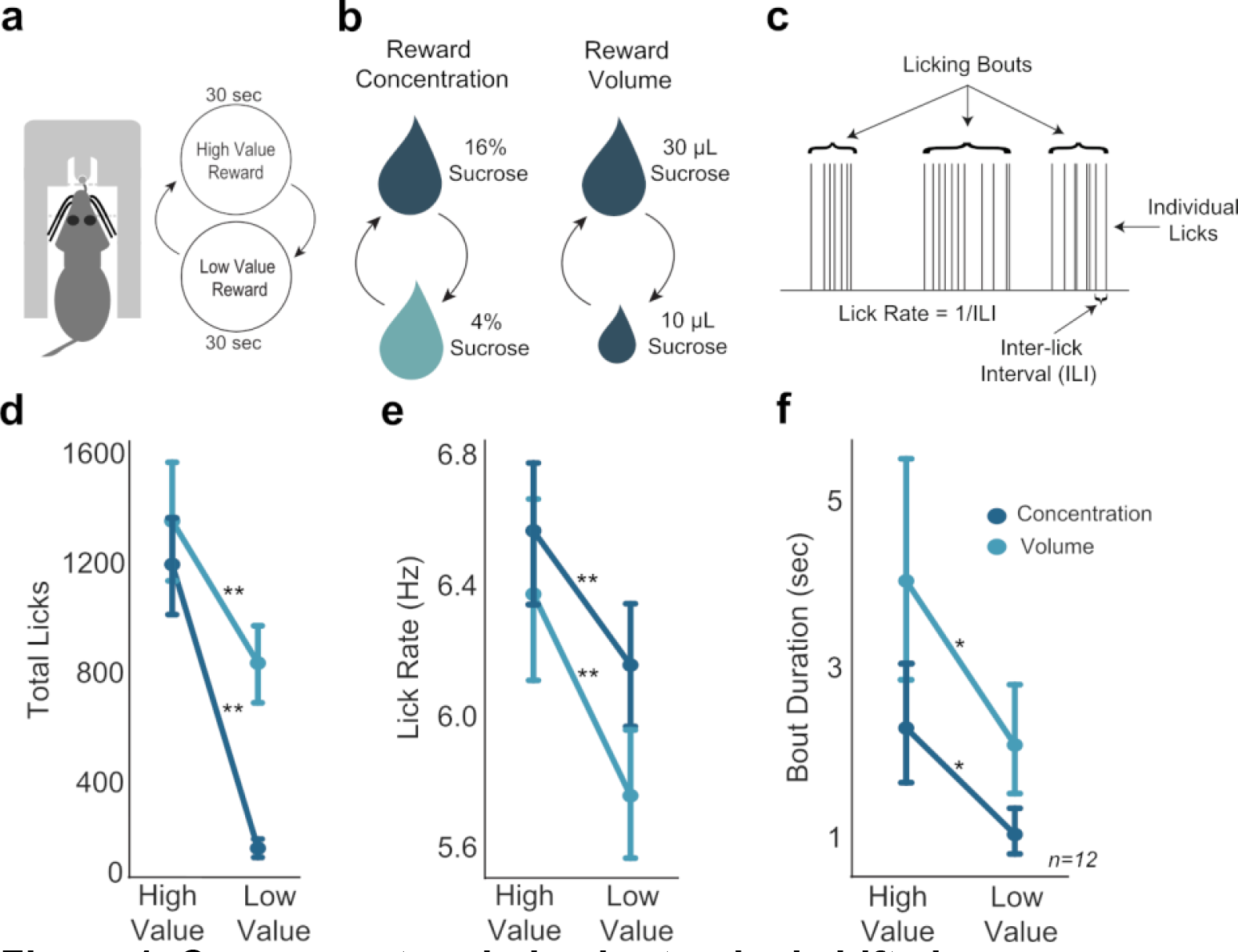
Consummatory behavior tracked shifts in sucrose concentration and fluid volume. A. In the Shifting Values Licking Task, rats received access to one of two values of reward, with rewards alternating every 30 sec. B. Manipulation of reward value by changing either concentration or volume. C. Types of behavioral licking measurements recorded in all licking tasks. D,E,F. Rats licked more (D), faster (E), and over longer bouts (F) for the high concentration and large volume rewards. Single asterisk (*) denotes p<0.05; Double asterisk (**) denotes p<0.001. Error bars represent 95% confidence intervals.

Several measures of licking behavior varied with sucrose concentration or fluid volume: lick counts, inter-lick intervals, lick rate, and bout duration (Figure 1C). All rats licked more for the high concentration reward compared to the low concentration reward (paired t-test; t(11)=10.76, p<0.001) (Figure 1D). Rats also licked at a faster rate for the high concentration reward compared to the low concentration reward (paired t-test; t(11)=6.347, p<0.001) (Figure 1E). Additionally, rats had increased bout durations when licking for the high concentration reward compared to the low concentration reward (paired t-test: t(11)=2.943, p=0.013) (Figure 1F). There was no difference in variability of high or low concentration licks: the coefficient of variation for inter-lick intervals was the same (paired t-test: t(9)=0.864, p=0.41).

Rats behaved similarly when consuming the high concentration and large volume rewards. In volume manipulation sessions, rats emitted more licks for the large reward than the small reward (paired t-test; t(11)=4.99, p<0.001). However, this difference in lick counts was less robust than the difference in high and low concentration rewards during concentration manipulation sessions (Figure 1D). Rats licked at a faster rate for large rewards compared to small volume rewards (paired t-test; t(11)=6.311, p<0.001) (Figure 1E), and licking bouts were longer for large rewards compared to bouts to consume small rewards (Figure 1F), (paired t-test; t(11)=2.569, p=0.027).

### Shifting Values Licking Task: Coherent fluctuations in the theta range in the MFC and OFC

LFPs in the MFC (N=56) and OFC (N=64) from 4 rats with arrays implanted in both cortical areas were recorded during the standard Shifting Values Licking Task. The LFPs were analyzed with cross-correlation and a spectral method called directed coherence to assess the extent of coordinated fluctuations between the cortical regions (Figure 2A). Data from all rats tested with both shifts in sucrose concentration and fluid volume were used for this analysis. One of the rats had 16 LFPs recorded in each area (256 pairs). Two rats had 14 LFPs in MFC and 16 in OFC (224 pairs). The fourth rat had 12 LFPs in MFC and 16 in OFC (192 pairs). Data from a total of 896 electrode pairs were analyzed. As shown in Figure 2B, LFPs from both areas showed frank fluctuations during periods of sustained licking (bouts). Standard (non-directional) coherence for the LFPs peaked around a value of 0.6 near the licking frequency (Figure 2C). By measuring cross-correlation over a range of lags (time-domain directionality), we found evidence for near zero-lag correlations. (This analysis is done in the time domain and does not account for frequency-specific directional influences.)

**Figure 2.**
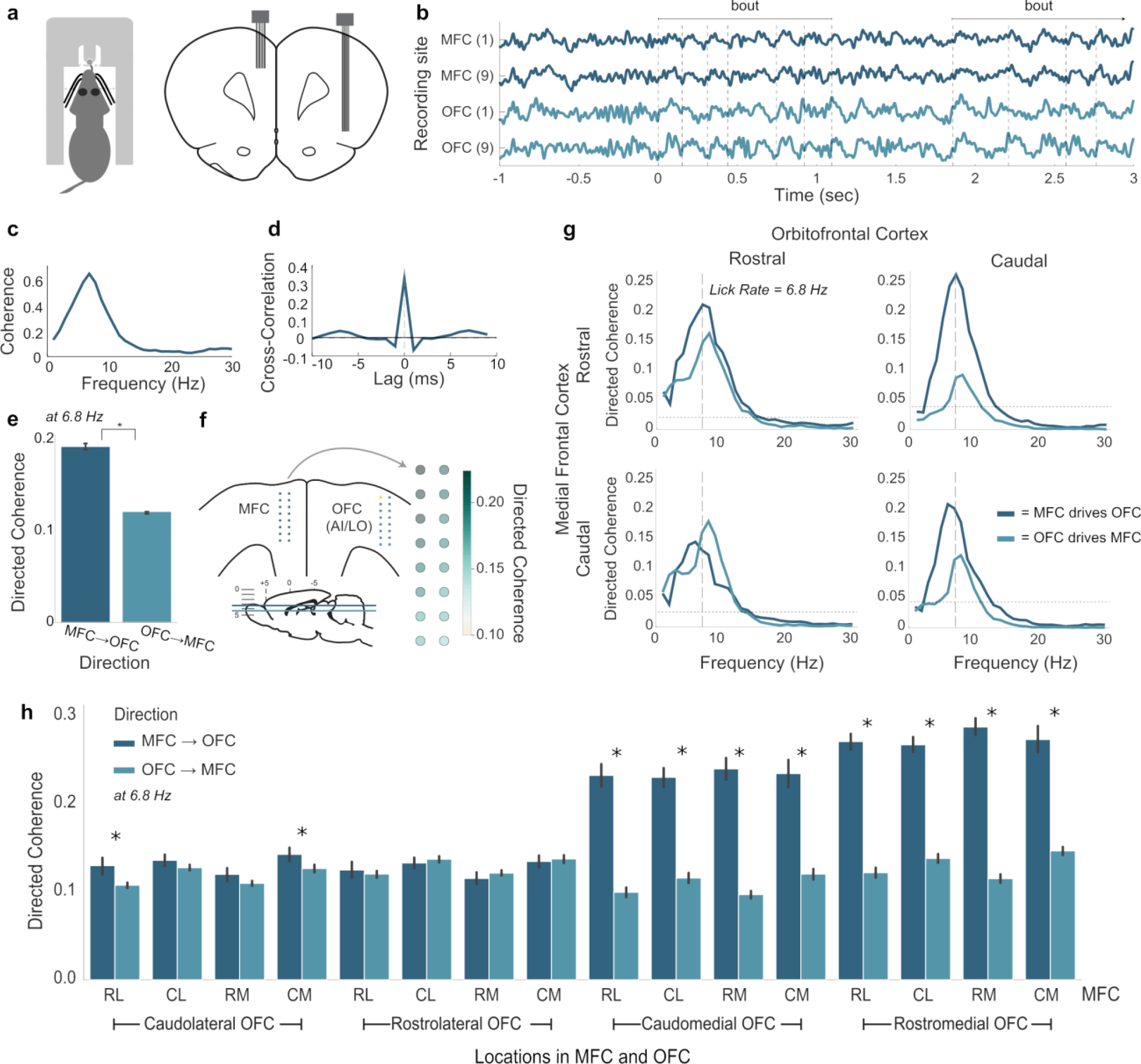
Coherent licking-related theta-band activity in the medial and orbital frontal cortices. A. Depiction of rat performing the Shifting Values Licking Task (left) and placement of recording arrays in the medial and lateral frontal cortices (right). B. Traces of simultaneous LFP recordings from rostral (1) and caudal (9) recording sites on arrays implanted in MFC and OFC. Two licking bouts are noted and the times of licks are shown as dashed vertical lines. C. Standard (non-directional) coherence between a pair of LFPs from MFC and OFC showed a peak near the licking frequency (∼7 Hz). D. Cross-correlation (time domain) showed a central peak with lag near 0 ms. E. Directed coherence at the licking frequency for MFC→OFC and OFC→MFC over all pairs of LFPs recorded in 4 rats. Asterisk denotes p<10^-6^ for effect of direction on coherence. F. Anatomical map of directed coherence values over one of the arrays. G. Directed coherence over frequencies up to 30 Hz, plotted for rostral and caudal sites in the MFC and OFC (panel B). H. Group summary of directed coherence over all pairs of recordings. Asterisks (*) denotes p<0.05. Error bars represent 95% confidence intervals.

Directed coherence values at the licking frequency were larger for MFC leading OFC compared to OFC leading MFC (Figure 2E). Notably, the magnitude of the coherence was no more than 0.2, suggesting a potential weak influence of MFC on the timing of fluctuations in OFC. The magnitude of the coherences were variable over electrodes, and plots of the measures onto the anatomical arrangement of the recording arrays revealed a gradient of directed coherence, with most rostral sites in MFC having larger coherence values compared to caudal sites (an example is shown in Figure 2F).

To further examine the role of spatial location on directed coherence, we denoted the locations of the recordings along the arrays as rostral or caudal (i.e. for each linear array with 8 electrodes, the four most rostral electrodes were denoted as rostral and the rest as caudal). An example of directed coherence over frequencies up to 30 Hz for the rostral and caudal sites (Figure 2B) is shown in Figure 2G. Directed coherence was larger for the direction MFC → OFC for most rostral electrode in the MFC and both the rostral and caudal electrodes in the OFC. The caudal electrode in the MFC had larger directed coherence for the direction MFC → OFC for the caudal, but not the rostral, electrode in OFC. A group summary of these findings, at the licking frequency, is shown in Figure 2H. Here, the locations of the electrodes was further split as medial and lateral, and differences in directed coherence were apparent for rostral and caudal sites in the MFC and medial sites in the OFC (right half of the plot). Directed coherence was equivocal for rostral and caudal sites in the MFC and lateral sites in the OFC (left half of the plot). Based on anatomical mapping of the arrays, the medial and lateral sites in the OFC were associated with the deep and superficial layers of the cortex, respectively. These findings suggest cross-laminar differences in the timing of the LFP fluctuations, with the rostral part of the MFC “driving” fluctuations in the deep layers of the OFC, and possibly serving as feedback from the MFC to the OFC (Gabbott et al., 2003).

### Shifting Values Licking Task: Lick entrainment in MFC and OFC tracks reward magnitude

We next aimed to determine if there were electrophysiological differences in MFC and OFC during access to the different types of rewards. Lick-field coherence (using methods originally developed for spike-field coherence in the Neurospec library for Matlab, Halliday et al., 1995). LFPs from both areas were coherent with licks at the licking frequency, and not at higher harmonic frequencies of licking (Figure 3A). Coherence levels were higher for licks that delivered high value fluid (concentration and volume) compared to low value fluid in MFC (paired t-test; Concentration: t(95)=39.972, p<0.001; Volume: t(95)=11.643, p<0.001) and OFC(paired t-test: Concentration: t(91)=17.386, p<0.001; Volume: t(91)=18.970, p<0.001) (Figure 3A-B). Furthermore, coherence was higher for high-value licks in the concentration shift sessions compared to the volume shift sessions in MFC (paired t-test; t(95)=6.901, p<0.001), but not the OFC (paired t-test; t(91)=-0.401, p=0.688). Phase angles at the licking frequency are shown in Figure 3C. With lick-field coherence ranging between 0 and 0.5, this analysis suggests that the LFP fluctuations at the licking frequency are only partially accounted for by the animals’ licking behavior and the extent of entrainment differs between cortical areas (larger in MFC) and is sensitive to reward value (larger for higher value fluid).

**Figure 3.**
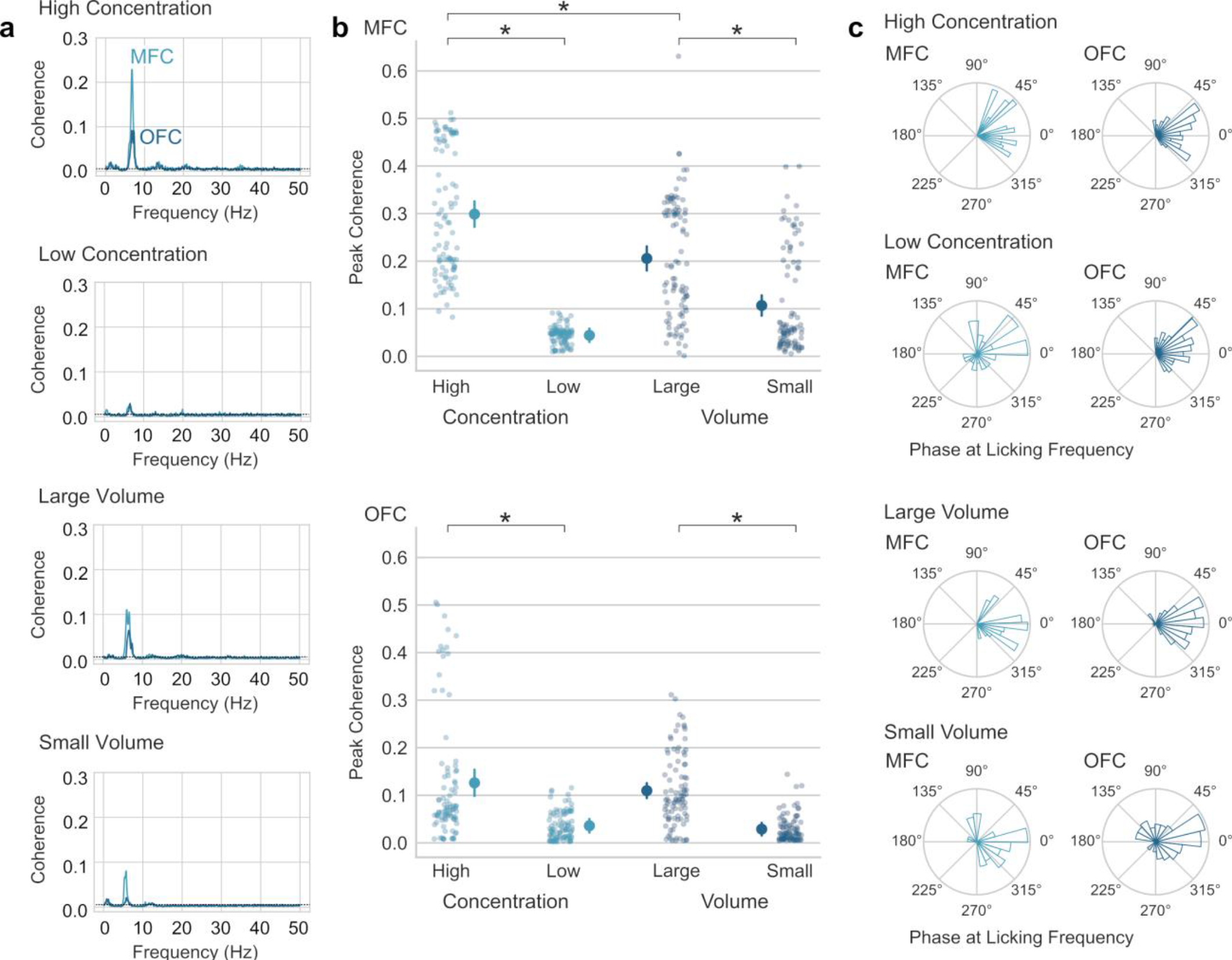
Lick-entrainment in the medial and orbital frontal cortices is sensitive to reward value. A. Average lick-field coherence for LFPs in MFC and OFC and licks for high and low concentration sucrose solutions and large and small fluid volumes. B. Peak coherence in the range of theta (4-12 Hz) for LFPs from MFC and OFC and licks for high and low concentration sucrose solutions and large and small fluid volumes. Asterisks (*) denotes p<0.05. Error bars represent 95% confidence intervals. C. Phase angles of LFPs at the licking frequency. Most LFPs were coherent with licks at phases between 45 and 315 degrees.

Three additional measurements of local field potential (LFP) activity were examined: amplitude (as measured by the size of Event-Related Potentials (ERP); Figure 4 - figure supplement 1A), spectral power (as measured by Event-Related Spectral Power (ERSP); Figure 4 - figure supplement 1B), and phase (as measured by Inter-Trial Coherence (ITC), Figure 4 - figure supplement 1C). Similar to results from lick-field coherence, we found lick-entrained activity in MFC and OFC that varied with both differences in sucrose concentration and fluid volume (Figure 4). Event-related potentials showed evidence for time-locked rhythmic fluctuations in LFPs from both cortical areas (Figure 4B,F). Both cortical areas showed elevated ITC between 4 and 8 Hz for licks that delivered the high concentration liquid sucrose but not the low concentration sucrose (Figure 4C,G). That is, the phase angles of the LFP fluctuations at the times of licks were more consistent when rats consumed the high concentration fluid compared to the low concentration fluid. This result was observed in all rats that were tested (dark blue lines in Figure 4D,H) (MFC: F(1,278)=443, p<0.001; OFC: F(1,177)=77.31, p<0.001; one-way ANOVAs with an error term for within-subject variation). Analysis of phase coherence (Figure 4 - figure supplement 1D) and event-related power (Figure 4 - figure supplement 1E) revealed effects solely in the 4-8 Hz (theta) frequency range.

**Figure 4.**
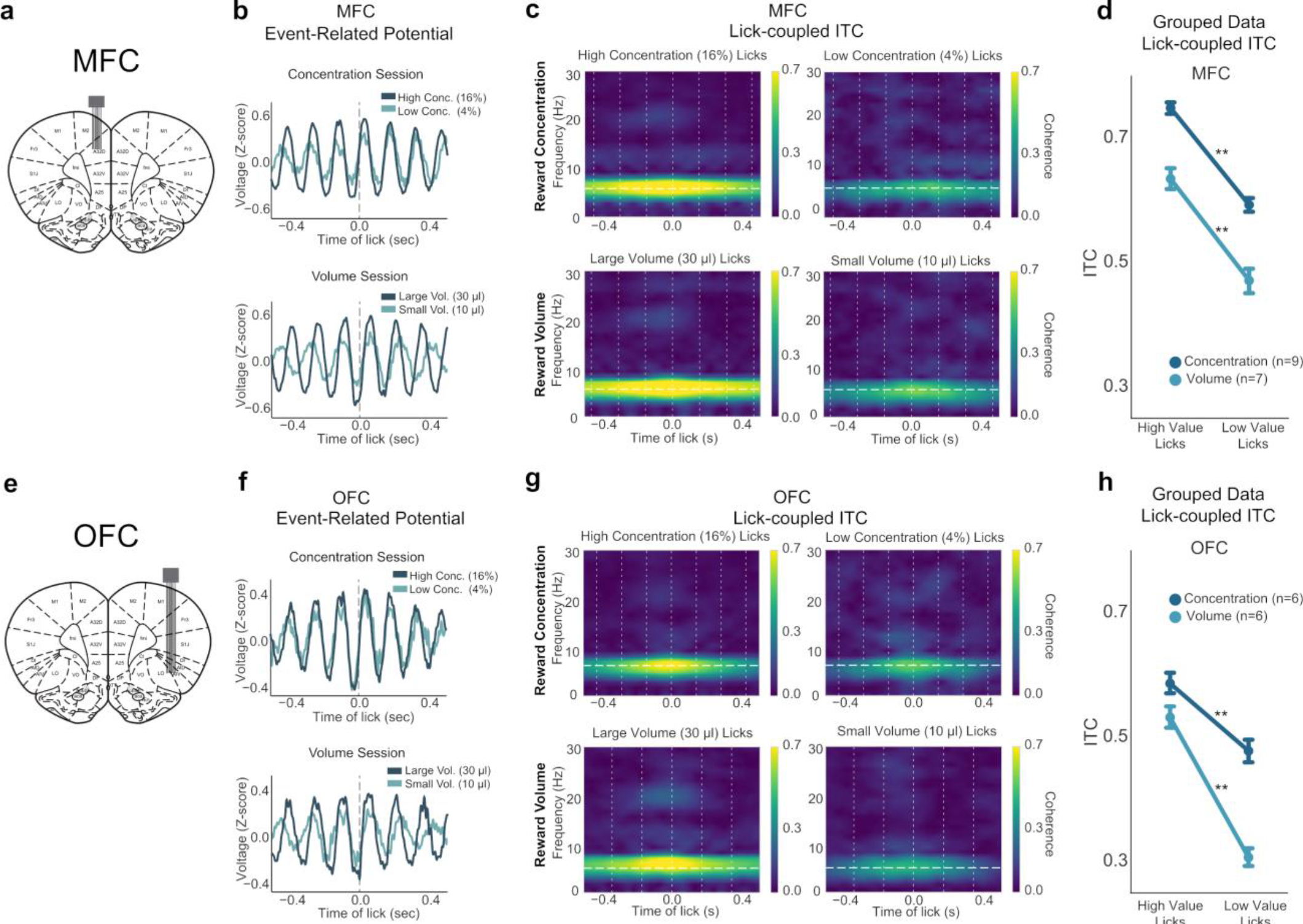
Lick-entrained neural activity in MFC and OFC tracked shifts in sucrose concentration and fluid volume. A,E. Rats were implanted with a 2x8 electrode array in either MFC (A) or OFC (E); representative coronal sections are shown. B,F. Event-related potentials during concentration and volume manipulation sessions in the Shifting Values Licking Task for MFC (B) and OFC (F). C,G. Spectral ITC time-frequency plots revealed strong phase locking during licks for the high concentration and large volume (left sides) rewards in both MFC (C) and OFC (G). Plots are from one electrode from one individual animal. ITC is consistently strongest around 4-8 Hz. D,H. Grouped data from all rats in both concentration and volume sessions in MFC (D) and OFC (H) showed strongest ITC during licks for the high value reward. Double asterisk (**) denotes p<0.001. Error bars represent 95% confidence intervals.

To assess differences in power, we used a peak-to-peak analysis of ERPs during licks for the high-value and low-value rewards. The measure calculates the difference in the maximum and minimum ERP amplitude using a window centered around each lick. The size of the window was twice each rat’s median inter-lick interval. LFPs in MFC showed increased amplitudes for high concentration rewards, as opposed to low concentration rewards (one-way ANOVA: F(1,278)=34.19, p<0.001). Figure 4B shows MFC ERPs for high and low concentration rewards of an example rat. This effect was not significant in OFC ERPs, as seen in Figure 4F (F(1,177)=0.557, p=0.456). We also measured ERSP, and although there was a decrease in MFC power from licks for the high to low concentration rewards specifically in the 4-8 Hz range (F(1,278)=18.72, p<0.001; one-way ANOVA), post-hoc testing revealed no relevant significance between high and low concentration licks (p=0.413). There was no major difference in ERSP measures in OFC (F(1,177)=0.039, p=0.843).

In sessions with shifts in fluid volume, event-related potentials in MFC or OFC did not distinguish between large versus small volume rewards (MFC: F(1,216)=0.865, p=0.354; OFC: (F(1,179)=1.876, p=0.173); one-way ANOVAs) (Figure 4B,F, bottom). There was no major difference in event-related spectral power during licks for large or small rewards in MFC or OFC (MFC: F(1,216)=0.877, p=0.35; OFC: F(1,179)=1.76, p=0.186); one-way ANOVAs). However, in both MFC and OFC, rats showed similar 4-8 Hz phase-locking for large rewards (Figure 4C,G, bottom), closely resembling what we observed with high concentration rewards (Figure 4C,G, top). Phase-locking was significantly increased for small rewards (MFC: F(1,216)=138.5, p<0.001; OFC: F(1,179)=280.8, p<0.001; one-way ANOVA) and was observedin all rats that were tested (light blue lines in Figure 4D,H).

These findings suggest that LFP activity in both MFC and OFC similarly encodes aspects of preferred versus less preferred reward options. 4-8 Hz phase-locking was strongest for both the high concentration and large volume rewards, which may be evidence that the animal is acting within a preferred state with the goal of obtaining their most “valued” reward. These findings provided further evidence suggesting that the entrainment of neural activity in MFC and OFC to the lick cycle tracks reward magnitude.

### Blocked-Interleaved Task: Engagement in and the vigor of licking vary with reward expectation

The same group of 12 rats were subsequently tested in an adjusted version of the Shifting Values Licking Task, which will be referred to as the Blocked-Interleaved Task (Figure 5A). In the first three minutes of the task, i.e. the “blocked” phase, rats behaviorally showed their typical differentiation of high versus low concentration rewards by emitting more licks for the high concentration reward (Figure 5B, left), and licked at a faster rate (Figure 5C, left). However, this pattern changed when the rewards were randomly presented in the “interleaved” part of the task. With a randomly interleaved reward presentation, rats licked nearly equally for high and low concentration rewards (Figure 5B, right; see also Figure 5 - figure supplement 1). We performed a two-way ANOVA on the number of licks by each lick type (high or low concentration) and portion of the task (blocked or interleaved). There was a significant interaction between concentration of reward and the blocked or interleaved portion of the task (F(1,33)=24.51, p<0.001). Post-hoc analyses revealed that while there was a significant difference in high and low concentration licks during the blocked portion (p<0.001), there was no difference between high and low concentration licks during the interleaved portion of the task (p=0.98). These findings suggest that shifting from blocked to interleaved presentations of the two rewards increased the animals’ engagement in licking for the lower value fluid.

**Figure 5.**
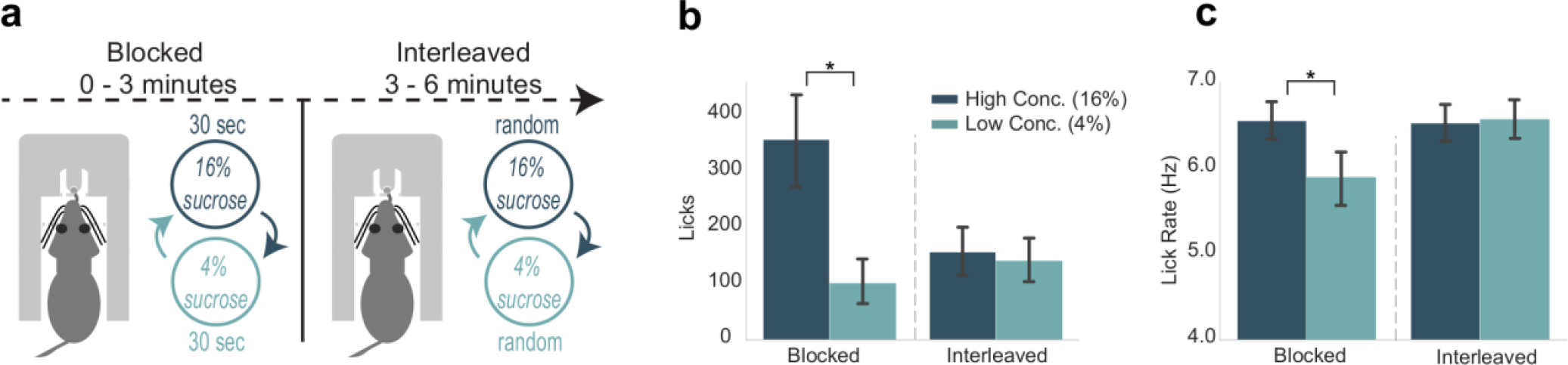
Engagement in and the vigor of licking varied with reward expectation. A. Rats participated in a modification of the Shifting Values Licking Task, called the Blocked- Interleaved Task, in which they received alternating access to high and low concentrations of liquid sucrose for three minutes and then received interleaved (and thus unpredictable) presentations of the two levels of sucrose for the rest of the session. B. Total licks emitted, a measure of task engagement, for both high and low concentration rewards during the blocked and interleaved portion of the task. Rats licked less for both rewards when rewards were randomly interleaved. C. Lick rate, a measure of response vigor, was similar for both rewards in the interleaved, but not blocked, portion of the task. Asterisk denotes p<0.05. Error bars represent 95% confidence intervals.

Additionally, there was a significant difference in lick rate by each lick type and portion of the task (F(1,33)=23.13, p<0.001; two-way ANOVA) (Figure 5C). Post hoc analyses revealed that rats licked significantly faster for high versus low concentration rewards during the blocked portion (p<0.005). Lick rates for high versus low concentration licks during the interleaved part of the task were not significantly different (p=0.99). Notably, lick rate during access to either high concentration (p=0.005) or low concentration (p=0.002) rewards during the interleaved portion was significantly increased from lick rate during access to the low concentration reward in the blocked portion of the task. These changes in lick rate were not accounted for by the changes in lick counts reported above (Spearman rank correlation: 0.44242, p=0.20042) and suggest that shifting from blocked to interleaved presentations of the two rewards increased the vigor with which the rats licked for the lower value fluid.

### Blocked-Interleaved Task: MFC rhythmicity tracks response vigor

Having established that the Blocked-Interleaved Task can reveal effects of reward expectation on task engagement and response vigor, we next examined how neural activity in the MFC and OFC varies with these behavioral measures. We assessed changes in lick-entrained ERPs and their amplitudes (Figure 6A,D), ERSP, and ITC (phase-locking) (Figure 6B- C,E-F). LFPs in MFC and OFC showed strong 4-8 Hz phase-locking during licks for the high concentration rewards in the blocked phase of the task (Figure 6B,E). We performed a two-way ANOVA on maximum ITC values (Figure 6C,F) from LFPs in both MFC and OFC for each rat and each electrode channel with interaction terms for lick type (high or low concentration reward) and portion of the task (blocked or interleaved reward access), and found a significant interaction of lick type by portion of the task (MFC: F(1,572)=10.45, p=0.001); OFC: F(1,363)=12.119, p<0.001). Post-hoc analyses revealed that while there was a significant difference in phase-locking of licks for high versus low concentration in the blocked portion (MFC: p<0.001; OFC: p<0.036), there was no difference in phase-locking of licks for high versus low concentration rewards in the interleaved portion of the task (MFC: p=0.999; OFC: p=0.973). In MFC, a two-way ANOVA revealed a significant interaction of lick type by portion of the task with ERP peak-to-peak size (Figure 6A) as the dependent variable (F(1,564)=6.232, p=0.013). However, there were no differences between the ERP measures between high and low concentration licks during the blocked portion of the task (p=0.887) and between high and low concentration licks during the interleaved portion of the task (p=0.938).

**Figure 6.**
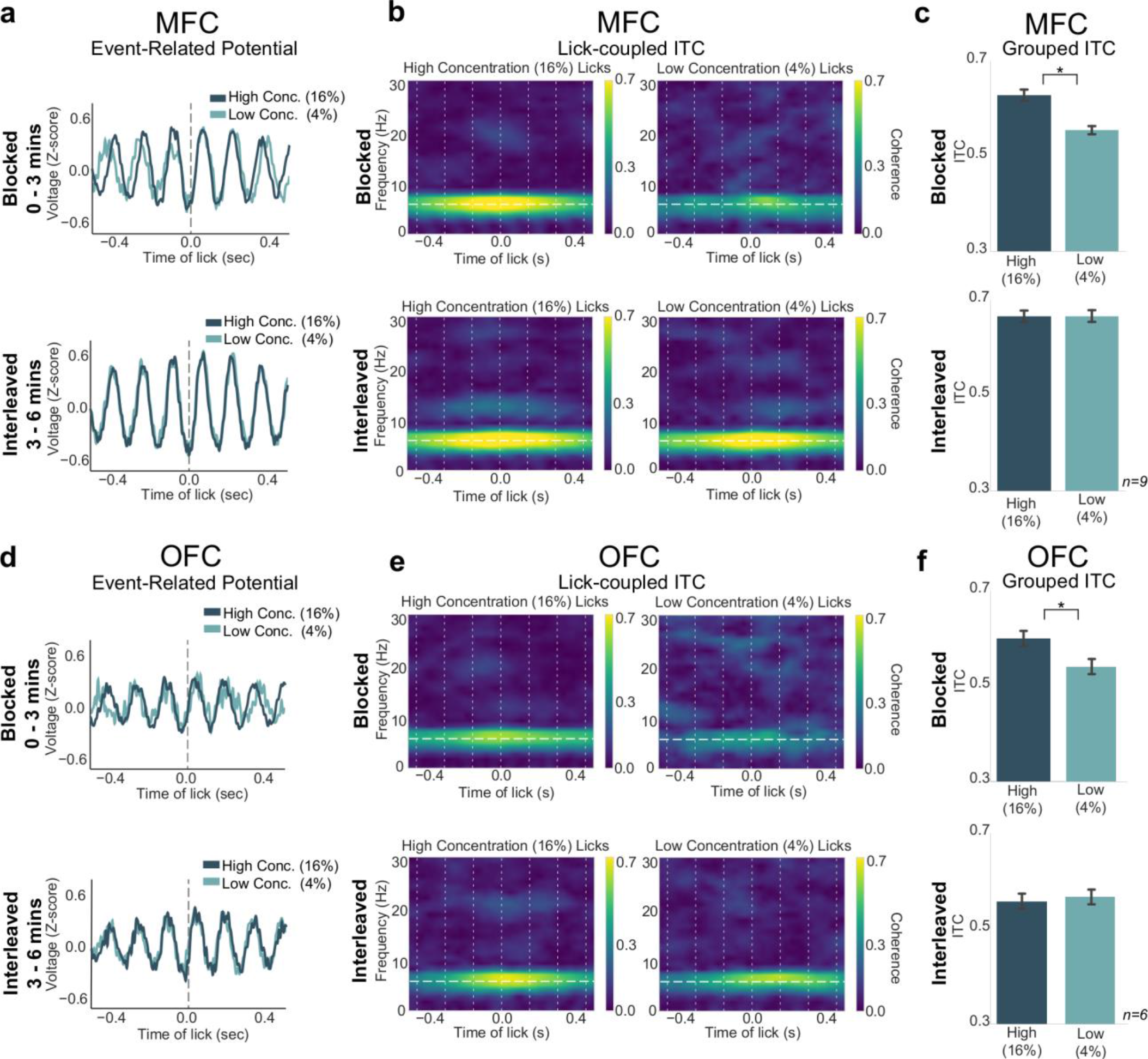
Lick-entrained neural activity varied with reward expectation. A,D. ERPs for licks of both rewards in MFC (A) and OFC (D) remain unchanged during the interleaved portion of the task. B,E. Spectral ITC plots revealed stronger 4-8 Hz phase-locking during licks for the high concentration reward in the blocked portion (top), but phase-locking during licks for high and low concentration rewards in the interleaved portion were indistinguishable from each other. C,F. Grouped data revealed no difference in ITC values during high or low concentration licks in the interleaved phase. Asterisk denotes p<0.05. Error bars represent 95% confidence intervals.

The same was true with ERSP measures for MFC LFPs; There was a significant interaction between lick type and portion of the task (F(1,564)=30.17, p<0.001; two-way ANOVA), but no significant difference between ERSP values between high and low concentration licks in the blocked (p=0.213) or interleaved (p=0.743) portions of the task. In OFC (Figure 6D), there was no significant interaction of lick type and portion of the task by the amplitude size of the lick’s ERPs (F(1,363)=0.131, p=0.718; two-way ANOVA), and no difference in OFC ERSP values of lick type by portion of the task either (F(1,363)=0.744, p=0.389; two-way ANOVA).

We wanted to further investigate potential differences in MFC and OFC in the Blocked- Interleaved Task, since initial results show a general increase of ITC values from MFC in the interleaved portion of the task and a general decrease in ITC values from OFC. This was of particular interest since MFC ITC values varied with the lick rate, which increased for both the high and low concentration licks in the interleaved portion of the task. We directly compared ITC values in both regions with lick rate and total lick counts (Figure 7).

**Figure 7.**
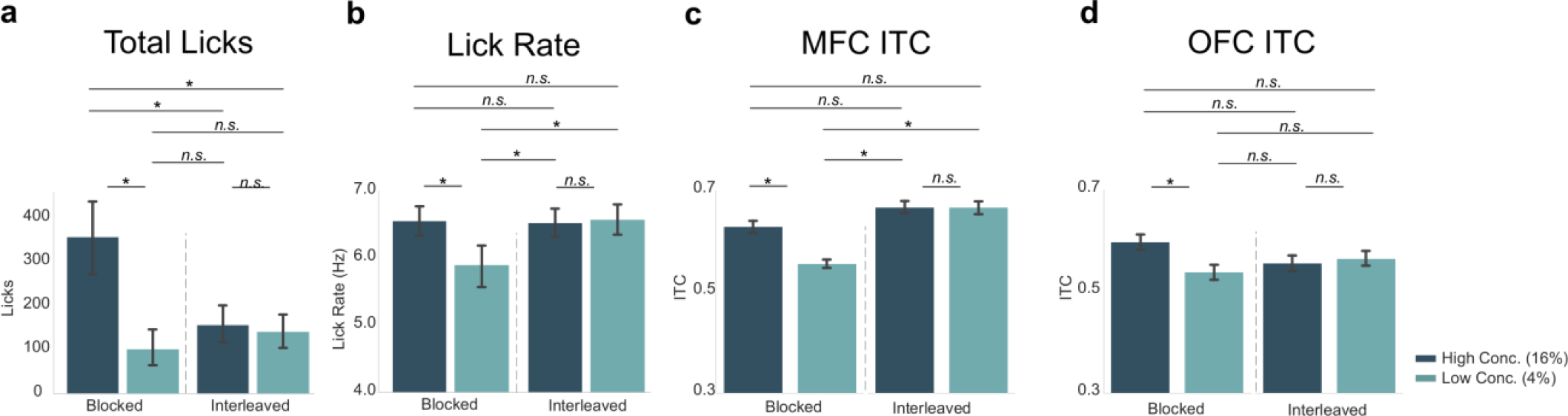
Neural activity in MFC, but not OFC, varied with the lick rate (vigor) and not task engagement (total licks). Post-hoc contrasts of statistically significant effects revealed by two-way ANOVA. Direct comparison of behavioral measures (A – total licks; B – lick rate) with MFC ITCs (C) and OFC ITCs (D) showed a similar pattern (and identical post-hoc statistical contrasts) between lick rate (B) and MFC ITCs (C). The pattern of post-hoc contrasts for OFC ITCs (D) did not match either total licks or lick rate. Asterisk denotes p<0.05. Error bars represent 95% confidence intervals.

Post-hoc analyses displayed in Figure 7C revealed that in MFC there was a significant difference between ITC values for the high versus low concentration licks (as also documented at the top of Figure 6C), but ITC values for high concentration licks during the blocked portion of the task did not differ from ITC values for either the high (p=0.075) or low concentration (p=0.089) conditions in the interleaved portion of the task. The pattern of post-hoc contrasts matches the lick-rate data (Figure 7B) for all paired comparisons. This match includes the finding (Figure 7C) that ITC values for low concentration licks in MFC differed from all three of the other conditions (high concentration blocked, high concentration interleaved, and low concentration interleaved licks; p<0.001 for each comparison). The MFC ITC post-hoc test results (Figure 7C) did not match the pattern for total licks (Figure 7A).

In OFC, ITC values (Figure 7D) did not match either the total-lick (Figure 7A) or lick-rate (Figure 7B) comparisons, despite the qualitative similarity with the total number of licks (compare Figure 7D with Figure 7A). The only significant difference in ITC values in OFC was between the high and low concentration licks in the blocked portion of the task (as also documented at the top of Figure 6F). All other comparisons were non-significant. This pattern of post-hoc comparisons did not match either total licks (compare Figure 5A with 5D) or lick rate (compare Figure 7B with 7D).

Together with the results summarized in Figure 6, these findings from post-hoc testing in Figure 7 provide evidence that MFC and OFC encode different aspects of licking and reward value. There was a clear match between the pattern of lick entrainment in the MFC, but not the OFC, with the animals’ licking rates. The correspondence between lick entrainment in MFC and the animals’ lick rates provides support for the idea that neural activity in MFC is sensitive to response vigor. By contrast, OFC might be involved in more general aspects of motivation, e.g. to lick or not (reward evaluation) based on reward magnitude or the predictability of the environment.

### Three Reward Task: Behavioral evidence for effects of relative reward value

The previous experiments assessed comparison of two levels of rewards (either high/low concentration or large/small volume) in the Shifting Values Licking Task. After finding behavioral and electrophysiological differences between two rewards, we aimed to investigate how animals process reward with contexts involving three different rewards. In this experiment, we assessed if rats process rewards in a relative manner or in an absolute manner by implementing a third intermediate (8% wt./vol. sucrose concentration) reward.

In the Three Reward Task (Figure 8A), the first block consists of the Shifting Values Licking Task with 30 sec shifts between the intermediate value (8% sucrose) reward and the low value (4% sucrose) reward. After 3 minutes the second block of the task begins, where rats then experience shifting values of reward from the high value (16% sucrose) reward to the intermediate value (8% sucrose) reward. This allowed us to assess how rats would process the intermediate 8% sucrose reward when it is paired with a worse (4%) or better (16%) option within one session. Additionally, the design introduces a second context (just like in the Blocked-Interleaved Task previously) in which we could assess if animals are still processing a (temporally) local comparison of reward types.

**Figure 8.**
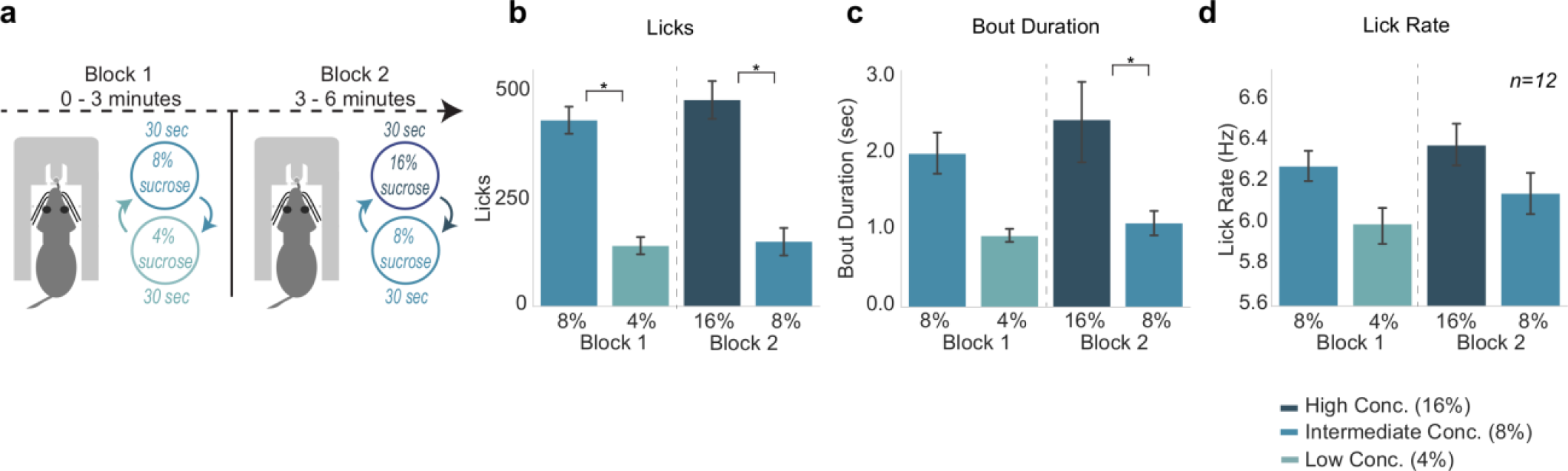
Consummatory behavior tracked relative differences in reward value. A. The Three Reward Task is a variation of Shifting Values Licking Task but with a third reward introduced. In the first block of the task, rats experience the intermediate (8%) reward and low (4%) reward. In block 2, rats experience the high (16%) reward paired with the intermediate (8%) reward. B. Rats licked more for the sweeter reward in each block. C. Rats showed greater bout durations for the sweeter reward. D. Lick rate showed a similar pattern to licks and bout duration, but was not statistically significant. Asterisk denotes p<0.05. Error bars represent 95% confidence intervals.

Licking varied with both reward value and block, i.e. low vs intermediate and intermediate vs high (F(3,33)=34.2, p<0.001) (Figure 8B). Post-hoc analyses revealed that rats emitted significantly more licks for the intermediate value 8% reward as opposed to the low value 4% reward in block 1 (p<0.001). In block 2, rats also emitted significantly fewer licks for the intermediate value 8% reward when it was paired with the high value 16% reward (p<0.001).

Rats also licked significantly less for the intermediate 8% reward in block 2 than they did in block 1 (p<0.001).

There was a more subtle effect for differences in bout duration across the different rewards (F(3,33)=5.333, p=0.004; two-way ANOVA) (Figure 8C). Post-hoc analyses revealed no significant difference in bout duration for the 4% versus 8% in block one (p=0.098), yet there was a significant decrease in bout durations during access to the 8% versus 16% in block two (p=0.023). Bout durations during access to the intermediate 8% reward in block 1 versus block 2 were not different (p=0.20). While there was a significant effect of lick type on lick rate (F(3,33)=10.59, p<0.001; two-way ANOVA), post-hoc analyses revealed no major differences in lick rate of the licks for rewards in block 1 (p=0.17) or block 2 (p=0.31) (Figure 8D),nor for the lick rate for 8% licks in block 1 versus block 2 (p=0.76).

### Three Reward Task: Neural activity does not reflect relative reward value encoding

The behavioral measures summarized above established that the Three Reward Task can reveal effects of relative value comparisons. We next analyzed electrophysiological signals from MFC and OFC (Figure 9) to determine if they tracked the animals’ behavior in the task, and might encode relative differences in value, or some other aspect of value, such as the absolute differences between the three rewards. We found a significant difference between ITC values for the three different rewards in both MFC and OFC (MFC: F(3,627)=154.4, p<0.001; OFC: F(3,363)=13.29, p<0.001; two-way ANOVAs). Tukey post-hoc analyses revealed a difference in ITC values between intermediate and low licks in block 1 (MFC: p<0.001; OFC: p=0.003), and a difference in ITCs between high and intermediate licks in block 2 for MFC only (MFC: p<0.005; OFC: p=0.313) (Figure 9B-C,E-F). There was no difference between ITC values from intermediate (8%) block 1 and intermediate block 2 licks in both regions (MFC: p=0.881; OFC: p=0.705).There was a significant difference between MFC ITC values for block 1 intermediate (8%) licks and block 2 high (16%) licks (p=0.028), as well as a significant difference between MFC ITC values for block 1 low (4%) licks and block 2 intermediate (8%) licks (p<0.001). Signals from the OFC did not differ across these conditions

**Figure 9.**
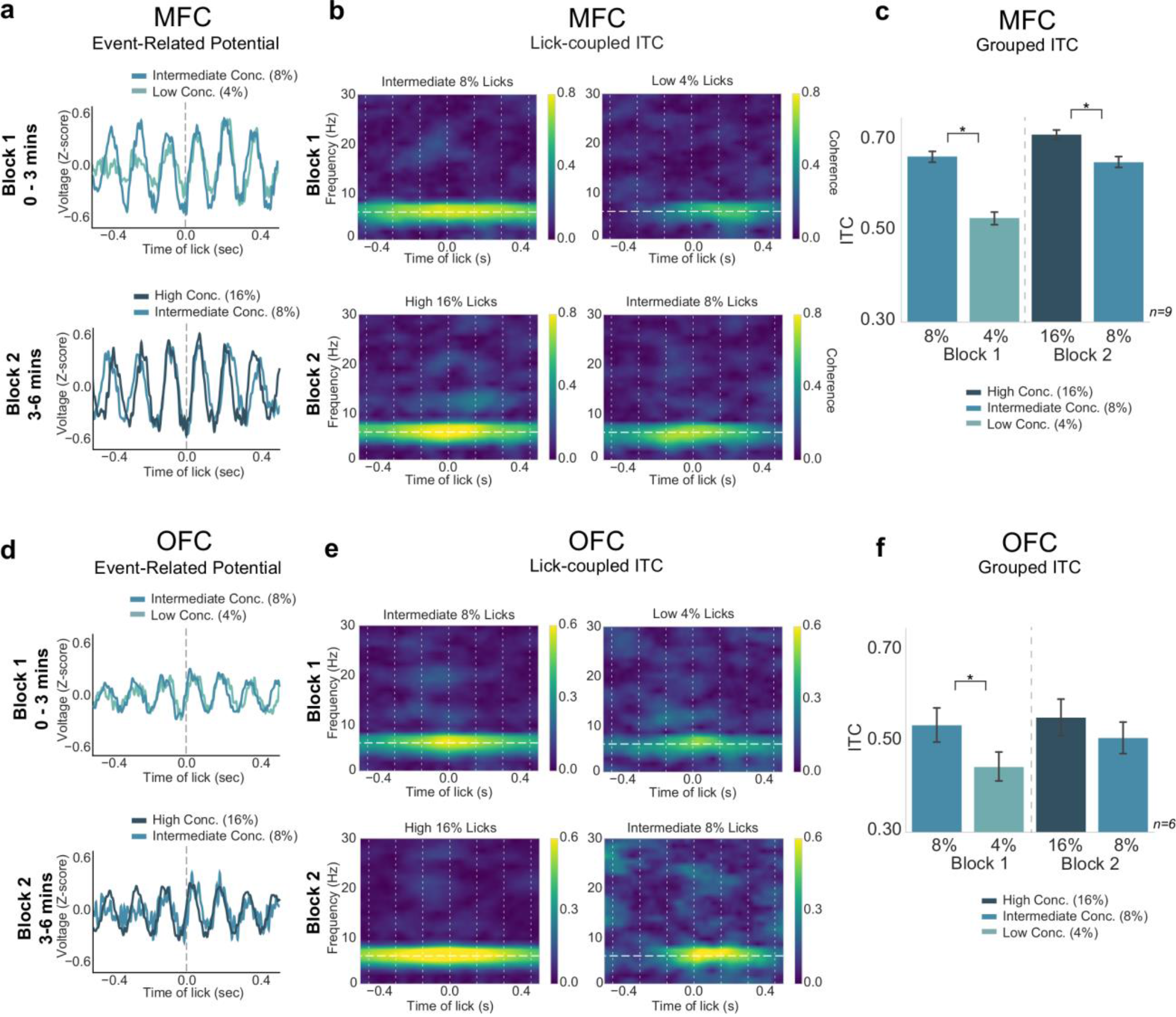
Neural activity in MFC, but not OFC, tracked absolute differences in reward value. A,D. ERPs for each block of the task from MFC (A) and OFC (D). B,E. ITC values in MFC (B) and OFC (E) showed strongest 4-8 Hz phase locking for the “high value” reward in each block. C,F. Group data revealed significantly greater ITC values for the high value reward in each block for MFC ITCs (C), and a similar pattern was found in OFC (F) but only block 1 rewards were significantly different. Asterisk denotes p<0.05. Error bars represent 95% confidence intervals.

Peak to peak amplitude analysis of the Three Reward Task revealed a significant effect of block on MFC LFP amplitude across lick types (F(3,627)=15.56, p<0.001; two-way ANOVA) (Figure 9A). Tukey post-hoc testing revealed no relevant significant differences between ERP size in MFC (between block 1 intermediate and low licks: p=0.864; between block 2 high and intermediate licks: p=0.944). There was no difference in OFC amplitude size (F(3,363)=0.827, p=0.479, two-way ANOVA) (Figure 9D). While there was a significant effect for ERSP values in both MFC and OFC (MFC: F(3,627)=18.35, p<0.001; OFC: F(3,363)=5.108, p=0.002; two-way ANOVAs), none of the relevant measures were significant (block 1 intermediate and low licks: MFC: p=0.875; OFC: p=0.492; block 2 high and intermediate licks: MFC: p=0.637; OFC: p=0.999).

The ITC findings, at least in MFC, support the idea that the “higher value” and “lower value” rewards in each context are being encoded differently across contexts. They indicate that MFC might instead encode absolute reward value instead of relative reward value. Qualitatively, the ITC values in MFC seem to have the same pattern as the lick rate (Figure 10B,C), similar to how MFC values reflected lick rate in the Blocked-Interleaved Task. However, post-hoc statistical testing revealed important differences. For example, the ITC in MFC differed significantly for high-vs. low-value rewards in both blocks 1 and 2, but lick rate did not. Importantly, post-hoc analyses revealed a significant difference in ITC values in MFC for every reward combination except for the intermediate block 1 and intermediate block 2 rewards, which reflects our operational definition for absolute encoding of value (see Figure 10 - figure supplement 1A-B).

**Figure 10.**
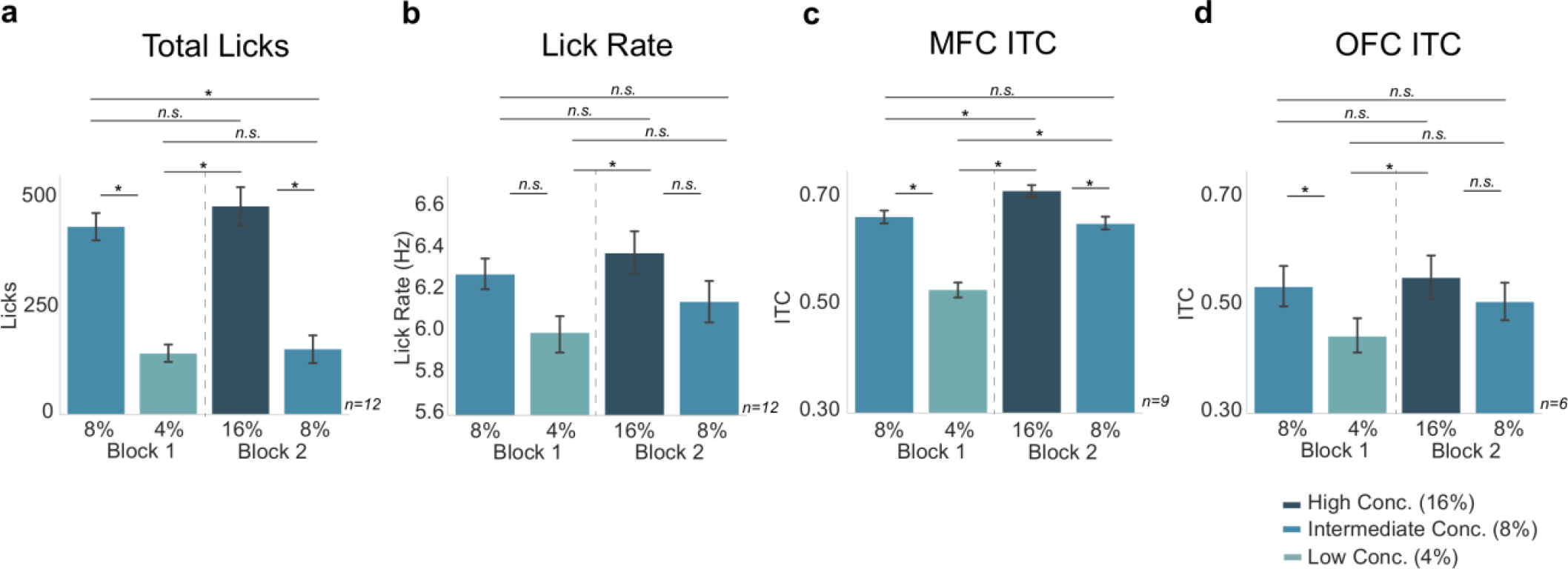
Neural activity in MFC, but not OFC, varied with effects of absolute reward value on lick rate (vigor) and task engagement (total licks). A,B. Behavioral measures replotted with significance bars for each combination reward. MFC ITCs (C) did not show the exact same pattern as lick rate, which is different from Figure 5. OFC ITCs (D) did not look like total licks or lick rate. Asterisk denotes p<0.05. Error bars represent 95% confidence intervals.

The encoding of value was less clear based on ITC measures from the OFC. These values did not directly match the licking behavior (in either rate, total licks, or bout duration) (compare Figure 10A,B with 8D), and did not show clear evidence for either absolute or relative encoding of reward. Instead, the results from Figure 10D indicate that OFC might instead encode reward value in a mixed absolute/relative manner (as in Figure 10 - figure supplement 1C and Figure 10 - figure supplement 2). However, these findings should be interpreted in the light of uneven sampling between areas, with fewer recordings done in the OFC. It is therefore possible that our results are underpowered for the OFC and new experiments could reveal an alternative interpretation.

## DISCUSSION

We investigated the role of MFC and OFC in processing reward information as rats participated in various consummatory licking tasks. Rats process and express changes in reward size in roughly the same manner as with reward concentration, both behaviorally and electrophysiologically. LFP activity in both MFC and OFC is sensitive to changes in reward type (both volume and concentration). Our results reveal context-dependent value signals in both regions through randomly presented rewards and by introducing a third reward in the task. Behaviorally, rats show evidence for a relative expression of rewards, while neural activity in MFC and OFC did not reflect relative encoding of reward. Together, our findings suggest that rats sample rewards and commit to consuming a given reward when they are able to predict its value, and this behavior is coupled to neural activity in MFC and OFC that encode both the value of the reward and the animal’s consummatory strategy. The subtle differences between the two regions follow the hypothesis that these areas provide different roles during consummatory behavior. We additionally provide evidence for MFC representing action-outcome relationships, as MFC ITC activity is more strongly correlated to the action of licking and may signal information about the “value of the action.”

### Rhythmic Activity and Reward Processing

Similar to our previous studies (Horst and Laubach, 2013; Amarante et al., 2017), neural activity was entrained to the lick cycle across all tasks in both MFC and OFC. Entrainment was strongest for the high-value reward (either of size or sweetness) and varied with the animals consummatory strategy (persistently lick a highly preferred option or sample fluid and wait for better option). Previous studies have viewed this rhythmic activity as being driven by the act of licking, as rats naturally lick at 6-7 Hz (Travers et al, 1997; Weijnen, 1998; Host and Laubach, 2013). However, the activity cannot be explained solely by licking, as there are instances where phase-locking and behavior do not show the same pattern (e.g. the Blocked-Interleaved experiment), and the variety of studies reported here and in Amarante et al. (2017) suggest a higher order role for rhythmic activity in the control of consummatory behavior.

Indeed, a major question for our study might be the extent to which lick-entrained oscillations in MFC and OFC can be dissociated from the act of licking. We adapted methods for spike-field coherence to examine this question (Figure 2). We found that the coherence between licks and rhythmic LFP signals in the licking frequency range was no larger than 0.5 (with lick-field coherence ranging between 0 and 1), was stronger for the MFC recordings compared to the OFC recordings and varied with the reward value of the consumed fluid. These findings suggest that the LFP oscillations are not simply driven by licking.

Furthermore, using directed coherence to examine directional influences of recordings in the two cortical areas, we found evidence for a weak directionality at the licking frequency such that the phase of the signals in MFC lead those in OFC. This result was especially apparent for the most rostral recording sites in the MFC, located in the medial orbital area and the adjacent frontal agranular area. Notably, these recording sites are immediately adjacent to a region of the frontal cortex where oral movements can be generated by electrical stimulation at low current (Yoshida et al., 2009). As such, the field recordings in the rostral part of the MFC might reflect activity from the adjacent oral motor cortex or could be locally generated. Resolving this matter will require new experiments, likely using optogenetic stimulation to avoid stimulation of fibers of passage.

We propose a functional interpretation of these signals based on findings on “medial frontal theta” (Cavanagh and Frank, 2014) in other types of behavioral tasks. There have been several proposals for the role of frontal theta in information processing. One idea is that the rhythm acts to break up sensory information into temporal chunks (Uchida and Mainen, 2003), and is related to the notion of a global oscillatory signal to synchronize neural activity across multiple brain structures throughout the taste-reward circuit (Gutierrez and Simon, 2013). Another idea is that frontal theta acts as an action monitoring signal (Cavanagh et al, 2012; Narayanan et al., 2013; Laubach et al., 2015), which can be generated through simple recurrent spiking network models (Bekolay et al., 2014). Finally, instead representing a specific function, frontal theta may act as a convenient “language” for distant brain regions to exchange information with each other (Womelsdorf et al., 2010). Our general findings contribute to this literature by suggesting that frontal theta acts as a value signal to guide consummatory behavior, which is the ultimate consequence of many goal-directed actions in natural environments.

### A Common Code for Reward Magnitude

A major finding in the present study (Figures 3 and 4) was the similar electrophysiological signals in MFC and OFC are associated with the consumption of high and low concentration liquid sucrose rewards and large and small volume rewards. Although other studies have found either decreases (Kaplan et al., 2001) or increases in behavior with increases in concentration and volume rewards in the same study (Hulse et al., 1960; Collier and Myers, 1961; Collier and Wills, 1961), these studies did not investigate the electrophysiological correlates of consuming rewards. Our study is the first to show a generalized “value signal” in the frontal cortex that scales with increased size and increased concentration of liquid sucrose. These signals might underlie the computation of a common currency (Montague and Berns, 2005; Levy and Glimcher, 2011; Levy and Glimcher, 2012; Strait et al., 2014) for the amount of nutrient available in a given food item and contribute to value-guided control of consumption.

### Evidence for the Contextual Control of Consumption

In the Blocked-Interleaved Task (Figure 5A), rats who licked more, longer, and faster for the high concentration reward when rewards were blocked did not continue to do so during interleaved portion of the task (Figure 5B-C). Instead, they licked nearly equally for the high and low concentration solutions, a result that is suggestive of the loss of positive contrast effects for the higher value fluid that is commonly found in the blocked design (Parent et al., 2015a). Despite these differences in behavior, the rats’ LFPs in MFC and OFC showed high levels of lick-entrained activity, essentially equal to that found during consumption of the higher value fluid in the blocked part of the session.

This finding is hard to reconcile with enhanced lick entrainment reflecting reward contrast effects. If positive contrast engenders entrainment, then LFPs should have shown reduced phase locking to the lick cycle in the interleaved portion of the task. Instead, the results might suggest that LFPs in MFC and OFC are entrained to licking when rats engage in persistent licking, as was found in the periods with high concentration access in the blocked part of the sessions and across the entire interleaved part of the session, and entrainment is reduced when rats switch to sampling the fluid during periods with low value access in the blocked part of the session. By this view, LFP entrainment to the lick cycle could serve as a contextual marker for reward state and the behavioral strategy deployed by the rat to sample and wait or persistently consume the liquid sucrose. This contextual information would depend on knowledge of the temporal structure of the reward deliveries. That is, when reward values are blocked, the rats have learned to expect alternative access to higher and lower reward values over extended periods of time (30 sec). By contrast, when reward values are interleaved, the changes in values occur rapidly and are unpredictable. The reduction in lick entrainment might therefore reflect the animal’s sampling strategy.

Contextual coding of reward value was also apparent in the Three Reward Task (Figures 8-10), where lick entrainment was stronger when the higher value option was available (Figure 9). In this case, the strength of engagement, for MFC but not OFC, tracked reward value in manner suggestive of an absolute reward encoding, with entrainment being higher for the 16% sucrose solution compared to the 8% solution when both were the “best” option (Figure 10C,D). These electrophysiological results were notably distinct from behavioral measures such as total licking output and lick rate (Figure 10A,B), which provided evidence for relative value comparisons.

Our electrophysiological results support theories of absolute reward value (Hull, 1943; Spence, 1956; Flaherty, 1982), as opposed to theories of relative reward value (Crespi, 1942; Black, 1968; Webber et al., 2015). Our findings might also fit with the neuro-economics idea of menu invariance versus menu-dependent goods (Padoa-Schioppa, 2011), both of which have been supported by electrophysiological studies on OFC (Padoa-Schioppa and Assad, 2006; Padoa-Schioppa and Assad, 2008; Tremblay and Schultz, 1999; Saez et al., 2017). Notably, in several instances we found a mismatch of behavioral output and corresponding magnitude of neural activity. This was evident in the Blocked-Interleaved task, where MFC and OFC ITCs did not reflect total licks emitted, as well as in the Three Reward Task where MFC and OFC ITCs did not reflect total licks or lick rate. This is in opposition to the Shifting Values Licking Task, where MFC and OFC activity directly matched behavioral output of licks, lick rate, and bout duration. These findings reveal the importance of recording careful behavioral output with electrophysiological recordings, and it remains an open discussion on the mechanisms behind correlative behavior versus diverging behavioral output from neural activity.

### Functional interpretations of phase entrainment

The original observation that suggested phase locking of licks to MFC field potentials was reported in Horst and Laubach (2013). Peri-event plots of LFPs around the times of licks revealed Event-Related Potentials (ERPs). The nature of ERPs has been researched extensively in the EEG literature. A leading view is that ERPs arise from a synchronization of the phase of an ongoing rhythm and/or from the superposition of inputs to the region of cortex near the electrode (e.g., Klimesch et al., 2007; Suaseng et al., 2007). Evidence for phase locking near the licking frequency can be found in Figures 1 and 3 in Amarante et al. (2017), with some exceptions being at slightly higher frequencies, e.g., Figure 8E in that study. By contrast, LFP power is typically tonic in the range of delta (<4 Hz) and the animals’ licking frequency mostly showed only minor changes in power (Figures 1 and 3 in Amarante et al., 2017). Furthermore, in another experiment with periodic reinforcement, we reported that phase but not power varied reinforcement (Figure 7, Amarante et al., 2017). These findings suggest that phase, not power, has a relationship with reinforced licking behavior, and the same determinants for phase locking likely apply to the results reported here.

Our data suggest that the act of licking synchronizes the phase of ongoing rhythms in the MFC and OFC and that this synchronization occurs during periods of sustained increases in delta band power. Computational models of LFP rhythmicity suggest that information flow is controlled by the interplay between different functional rhythms, with activity at higher frequencies nested within the periods of lower frequencies (Kopell et al., 2010). Brain slice (Carracedo et al., 2013) suggest that theta-range rhymicity may be nested within cycles of the lower frequency delta rhythm. For our studies, as shown in Figure 3 from Amarante et al. (2017), the duration of elevated phase synchronization was roughly twice as long as the median inter-lick interval, and the inverse of this interval would indicate a frequency of ∼3.5 Hz, i.e., delta. The same mechanisms are likely to apply to the present study. A possible source for the delta rhythm is the animal’s respiratory cycle (Lockmann and Tort, 2018), which must be regulated during periods of sustained licking when high value fluid is available.

Our finding of zero-lag correlation across frequencies (Figure 2D) further suggests that a source of common variance to MFC and OFC modulates the timing of processing, and leads to a slight advance in the phase angles of the rhythm in MFC relative to OFC (directed coherence in Figure 2E-H). The strength of this input would seem to vary between areas, being stronger in MFC compared to OFC and strongest in the rostral MFC (medial orbital cortex) overall (Figure 2H). These patterns of directional influences presumably vary with the extent of lick-field coherence and thereby with the value of the consumed fluid.

It is not clear from our studies if the reduction in entrainment when low value rewards are available is an active or passive process. For example, it is possible that some active input to the MFC and OFC denotes the temporal context (e.g. dopamine, hippocampus), enhancing entrainment when the higher value option is available. Alternatively, signals from sensorimotor regions of the frontal cortex, which sit in between the MFC and OFC, the oral sensory and motor cortices (Yoshida et al., 2009), might be reduced during periods with less intense licking, leading to a passive reduction in overall frontal lick entrainment. Future studies are needed to address these neural mechanisms of licking-related synchrony in the rodent frontal cortex.

### Differences in reward signaling between MFC and OFC

The electrophysiological results from the Blocked-Interleaved Task and Three Reward Task suggest that MFC and OFC, while showing similar results overall, may be contributing to processing reward information in different ways. It is important to note that due to a smaller sample size of OFC recordings, the less clear findings in OFC may indeed require further future experiments. However, our findings do follow previous work on subtle differences of these areas. In accord with a previous theory on proposed MFC and OFC functions (Balleine and Dickinson, 1998, Balleine and Dickinson, 2000; Schoenbaum et al., 2009; Sul et al., 2011; Passingham and Wise, 2012), MFC activity may be acting to maintain and optimize licking behavior in an action-centric manner, as reflected in measures such as the licking rate, a measure associated with vigor and sensitive to inactivation of the same cortical area in a progressive ratio licking task (Swanson et al., 2019). By contrast, OFC activity generally reflected differences in reward value, perhaps due to the different sensory properties of the fluids (Gutierrez et al., 2006), and was not sensitive to licking rate (vigor) or task engagement (total licks).

## METHODS

All procedures carried out in this set of experiments were approved by the Animal Care and Use Committee at American University (Washington, DC). Procedures conformed to the standards of the National Institutes of Health Guide for the Care and Use of Laboratory Animals. All efforts were taken to minimize the number of animals used and to reduce pain and suffering.

### Animals

Male Long Evans and Sprague Dawley rats weighing between 300 and 325 g were used in these studies (Charles River, Envigo). As relatively few animals were used, we did not investigate sex differences in reward processing in this study. Sex differences among rats are well known for how liquid sucrose is consumed (e.g. Sclafani et al., 1987) and classic studies of incentive contrast (e.g. Flaherty & Rowan, 1986), which led to the design of the behavioral procedures used here, were mostly carried out using male rats. As such, we cannot comment on sex differences or how reward value is encoded in the frontal cortex of female rats. These important topics require further study.

Key Resources Table (animals and any item with an RRID)

| Key Resources Table (animals and any item with an RRID) |  |  |  |  |
| --- | --- | --- | --- | --- |
| Reagent type (species) or resource | Designation | Source or reference | Identifiers (RRID) | Additional information |
| Strain, strain background (Rattus norvegicus) | Sprague-Dawley, Long-Evans | Charles River, Envigo | NA | Rat (male) |
| Instrument | Precision Syringe Pump Controller | https://doi.org/10.1523/ENEURO.0240-19.2019 | SCR_021493 |  |
| Software, Algorithm | Med-PC | MedAssociates | SCR_012156 |  |
| Software, Algorithm | GNU Octave | https://www.gnu.org/software/octave/ | SCR_014398 |  |
| Software, Algorithm | R Project for Statistical Computing | https://www.r-project.org/ | SCR_001905 |  |
| Software, Algorithm | NeuroExplorer | https://www.neuroexplorer.com/ | SCR_001818 |  |
| Software, Algorithm | Matplotlib | https://matplotlib.org/ | SCR_008624 |  |
| Software, Algorithm | IPython | https://ipython.org/ | SCR_001658 |  |
| Software, Algorithm | Jupyter | https://jupyter.org/ | SCR_018416 |  |
| Software, Algorithm | Seaborn | https://seaborn.pydata.org/ | SCR_018132 |  |
| Software, Algorithm | MATLAB | Mathworks | SCR_001622 |  |
| Software, Algorithm | EEGLab | https://sccn.ucsd.edu/eeglab/index.php | SCR_007292 |  |

Rats were given one week to acclimate with daily handling prior to behavioral training and surgery and were then kept with regulated access to food to maintain 90% of their free- feeding body weight. They were given ∼18 g of standard rat chow each day in the evenings following experiments. Rats were single housed in their home cages in a 12h light/dark cycle colony room, with experiments occurring during the light cycle. A total of 12 rats had a 2x8 microwire array implanted into either the MFC (N=6), the OFC (N=2) or one array in each area contralaterally (N=4). Arrays consisted of 16 blunt-cut 50-µm tungsten (Tucker-Davis Technologies) or stainless steel (Microprobes) wires, separated by 250 µm within each row and 500 µm between rows. In vitro impedances for the microwires were ∼150 kΩ.

### Surgeries

Animals had full access to food and water in the days prior to surgery. Stereotaxic surgery was performed using standard methods. Briefly, animals were lightly anesthetized with isoflurane (2.5% for ∼2 minutes), and were then injected intraperitoneally with ketamine (100mg/kg) and dexdomitor (0.25mg/kg) to maintain a surgical plane of anesthesia. The skull was exposed, and craniotomies were made above the implant locations. Microwire arrays were lowered into MFC (coordinates from bregma (AP: +3.2 mm; ML: + 1.0 mm; DV: -1.2 mm from the surface of the brain, at a 12° posterior angle; Paxinos and Watson, 2013) or into OFC (AP: +3.2 mm, ML: + 4.0 mm, DV: -4.0 mm; Paxinos and Watson, 2013). The part of the MFC studied here is also called “medial prefrontal cortex” in many rodent studies and the region is thought to be homologous to the rostral ACC of primates (Laubach et al., 2018). Four skull screws were placed along the edges of the skull and a ground wire was secured in the intracranial space above the posterior cerebral cortex. Electrode arrays were connected to a headstage cable and modified Plexon preamplifier during surgery, and recordings were made to assess neural activity during array placement. Craniotomies were sealed using cyanocrylate (Slo-Zap) and an accelerator (Zip Kicker), and methyl methacrylate dental cement (AM Systems) was applied and affixed to the skull via the skull screws. Animals were given a reversal agent for dexdomitor (Antisedan, s.c. 0.25 mg/ml), and Carprofen (5 mg/kg, s.c.) was administered for postoperative analgesia. Animals recovered from surgery in their home cages for at least one week with full food and water, and were weighed and monitored daily for one week after surgery.

### Behavioral Apparatus

Rats were trained in operant chambers housed within a sound-attenuating external chamber (Med Associates; St. Albans, VT). Operant chambers contained a custom-made glass drinking spout that was connected to multiple fluid lines allowing for multiple fluids to be consumed at the same location. The spout was centered on one side of the operant chamber wall at a height of 6.5 cm from the chamber floor. Tygon tubing connected to the back of the drinking spout administered the fluid from a 60-cc syringe hooked up to either a PHM-100 pump (Med Associates) for standard experiments, or to a customized open-source syringe pump controller (Amarante et al., 2019) that is programmed by a teensy microcontroller to deliver different volumes of fluid with the same delivery time from one central syringe pump. A “light-pipe” lickometer (Med Associates) detected licks via an LED photobeam, and each lick triggered the pump to deliver roughly 30 μL per 0.5 second. Behavioral protocols were run though Med-PC version IV (Med Associates), and behavioral data was sent via TTL pulses from the Med-PC software to the Plexon recording system.

### Shifting Values Licking Task

The operant licking task used here is similar to those previously described (Parent et al., 2015a,b; Amarante et al., 2017). Briefly, rats were placed in the operant chamber for thirty minutes, where they were solely required to lick at the drinking spout to obtain a liquid sucrose reward. Licks to the light-pipe lickometer would trigger the syringe pump to deliver liquid sucrose over 0.5 sec. In other words, the first lick to the spout triggers the pump and reward is then delivered for 0.5 sec, where any lick within that 0.5 sec window would also be rewarded. The next lick after 0.5 sec would subsequently trigger the pump to turn on again for 0.5 sec. Every 30 sec, the reward alternated between high (16% weight per volume) and low (4% wt./vol.) concentrations of liquid sucrose, delivered in a volume of 30 μL. In volume manipulation sessions, the reward alternated between a large (27.85 μL) and small volume (9.28 μL) of 16% liquid sucrose. Rewards were delivered over a period of 0.5 sec for all levels of concentration and volume using a custom-made syringe pump (Amarante et al., 2019). The animal’s licking behavior was constantly recorded throughout the test sessions.

### Blocked versus Randomly Interleaved Licking Task

The Shifting Values Licking Task was altered to allow for comparison of blocked versus interleaved presentations of reward values. The first three minutes of the task consisted of the standard Shifting Values Licking Task, with 30 second blocks of either the high or low concentration sucrose rewards delivered exclusively during the block. After three minutes, the rewards were presented in a pseudo-random order (e.g., high, high, low, high, low, low, high) for the rest of the test session. With rewards interleaved, rats were unaware of which reward would be delivered next. Behavioral and neural data were only analyzed from the first six minutes of each test session. We focused on manipulating sucrose concentration, and not fluid volume, in this task variation, as concentration differences provided the most effects of reward value on licking behavior (see Figure 1D).

### Three Reward Licking Task

The Shifting Values Licking Task was modified, using a third intermediate concentration of sucrose (8% wt./vol) to assess if reward value influenced behavior and neuronal activity in a relative or absolute manner. In the first three minutes of each session, rats received either the intermediate (8%) or low (4%) concentration of sucrose, with the two rewards delivered over alternating 30 second periods as in the SVLT. After three minutes, the rewards switched to the high (16%) and intermediate (8%) concentrations, and alternated between those concentrations for the rest of the session. Behavioral and neural data were only analyzed from the first six minutes of each test session.

### Electrophysiological Recordings

Electrophysiological recordings were made using a Plexon Multichannel Acquisition Processor (MAP; Plexon; Dallas, TX). Local field potentials were sampled on all electrodes and recorded continuously throughout the behavioral testing sessions using the Plexon system via National Instruments A/D card (PCI-DIO-32HS). The sampling rate was 1 kHz. The head-stage filters (Plexon) were at 0.5 Hz and 5.9 kHz. Electrodes with unstable signals or prominent peaks at 60 Hz in plots of power spectral density were excluded from quantitative analysis.

### Histology

After all experiments were completed, rats were deeply anesthetized via an intraperitoneal injection of Euthasol (100mg/kg) and then transcardially perfused using 4% paraformaldehyde in phosphate-buffered saline. Brains were cryoprotected with a 20% sucrose and 10% glycerol mixture and then sectioned horizontally on a freezing microtome. The slices were mounted on gelatin-subbed slides and stained for Nissl substance with thionin.

### Data Analysis: Software and Statistics

All data were analyzed using GNU Octave (https://www.gnu.org/software/octave/), Python (Anaconda distribution: https://www.continuum.io/), and R (https://www.r-project.org/). Analyses were run as Jupyter notebooks (http://jupyter.org/). Computer code used in this study is available upon request from the corresponding author.

Statistical testing was performed in R. Paired t-tests were used throughout the study and one or two-way ANOVA (with the error term due to subject) were used to compare data for both behavior and electrophysiological measures (maximum power and maximum inter-trial phase coherence) for high and low value licks, blocked versus interleaved licks, and high-intermediate-low licks. For significant ANOVAs, the error term was removed and Tukey’s post-hoc tests were performed on significant interaction terms for multiple comparisons. Descriptive statistics are reported as mean ± SEM, unless noted otherwise.

### Data Analysis: Behavior

All rats were first run for at least five standard sessions in the standard Shifting Values Licking Task with differences in concentration (16% and 4% wt./vol.). Rats have been shown to acquire incentive contrast effects in the SVLT after this duration of training (Parent et al., 2015a). For the Blocked-Interleaved and Three Reward tasks, rats were tested after extensive experience in the SVLT and after two “training” sessions with the Blocked-Interleaved and Three Reward designs. The electrophysiological recordings reported here were from the animals’ third session in each task.

Behavioral measures included total licks across the session, the duration and number of licking bouts, and the median inter-lick intervals (inverse of licking frequency). Bouts of licks were defined as having at least 3 licks within 300 ms and with an inter-bout interval of 0.5 sec or longer. Bouts were not analyzed in the Blocked-Interleaved Task; due to the unique structure of the task, bouts were all shortened by default due to a constantly changing reward in the interleaved phase of the task. While bouts of licks were reported in most tasks, electrophysiological correlates around bouts were not analyzed because there were often too few bouts (specifically for the low-lick conditions) in each session to deduce any electrophysiological effects of reward value on bout-related activity.

For analyzing lick rate, inter-lick intervals during the different types of rewards were obtained, and then the inverse of the median inter-lick interval provided the average lick rate in Hertz. Any inter-lick interval greater than 1 sec or less than 0.09 sec was excluded from the analysis. For licks during the randomly interleaved portion of the Blocked-Interleaved Task, more than two licks in a row were needed to calculate lick rate. To analyze behavioral variability of licks, we used coefficient of variation (ratio of the standard deviation to the mean) on high and low value inter-lick intervals that occurred within bouts.

In some experiments, imbalances were apparent in measures of total licks and lick rate. This was due in part to our only calculating inter-lick intervals that were less than 1 second and consisted of runs of at least 2 consecutive licks. As a result, some licks that were detected were not included in the quantitative measures of lick rate (e.g. two licks that occur 15 seconds apart from each other). Isolated licks occur in the behavioral design used in our studies when rats sample fluid from the spout during periods when low value fluid is available and then do not engage in persistent licking.

Total licks and lick rate are therefore distinct measures and will not always be coupled, especially because licks occur in bursts. Rats strongly engage when the higher value fluid is available in the blocked condition and alternatively will lick more sporadically and will default to sampling the fluid and not maintain engagement when the low value fluid is available. However, the rate of the licks, in said bouts or bursts, was higher overall during the interleaved parts of the tests sessions. Why this happened is not clear, but one interpretation is that rats are not suppressing or sampling the options anymore in the interleaved portion but are instead maintaining engagement in the task during the interleaved portion of the task when reward identity is unpredictable.

### Data Analysis: Local Field Potentials

Electrophysiological data were first analyzed in NeuroExplorer (http://www.neuroexplorer.com/), to check for artifacts and spectral integrity. Subsequent processing was done using signal processing routines in GNU Octave. Analysis of Local Field Potentials (LFP) used functions from the EEGLab toolbox (Delorme and Makeig, 2004) (Event-Related Spectral Power and Inter-Trial Phase Coherence) and the signal processing toolbox in GNU Octave (the peak2peak function was used to measure event-related amplitude). Circular statistics were calculated using the circular library for R. Graphical plots of data were made using the matplotlib and seaborn library for Python. Analyses were typically conducted in Jupyter notebooks, and interactions between Python, R, and Octave were implemented using the rpy2 and oct2py libraries for Python.

To measure the amplitude and phase of LFP in the frequency range of licking, LFPs were bandpass-filtered using eeglab’s eegfilt function, with a fir1 filter (Widmann and Schröger, 2012), centered at the rat’s licking frequency (licking frequency + inter-quartile range; typically around 4 to 9 Hz), and were subsequently z-scored.

Lick-Field Coherence (LFC) used routines (e.g. sp2a_m) from the Neurospec 2.0 library (http://www.neurospec.org/) for Matlab and GNU Octave. LFPs were low-pass filtered (100 Hz) using eegfilt.m from EEGLab. Directed Coherence also used routines (e.g. sp2a2_R2.m) from Neurospec 2.0. LFPs were low-pass filtered (100 Hz) using eegfilt.m from EEGLab. The following parameters were used for LFC and Directed Coherence: Segment power = 10 (1024 points, frequency resolution: 0.977 Hz), Hanning filtering with 50% tapering, and line noise removal for the LFPs at 60 Hz. Analyses focused on frequencies below 30 Hz based assessments of power spectra computed by the Neurospec library as part of this analysis. For inter-trial phase coherence (ITC) and event-related spectral power (ERSP), LFP data was preprocessed using eeglab’s eegfilt function with a fir1 filter and was bandpass filtered from 0 to 100 Hz. For group summaries, ITC and ERSP matrices were z-scored for that given rat after bandpass filtering the data. Peri-lick matrices were then formed by using a pre/post window of 2 seconds on each side, and the newtimef function from the eeglab toolbox was used to generate the time-frequency matrices for ITC and ERSP up to 30 Hz.

Since most of the lick counts from the Shifting Values Licking Task are generally imbalanced (with a greater number of licks for high versus low value rewards), we used permutation testing to perform analyses on amplitude and phase-locking in these studies. Licks were typically downsampled to match the lower number of licks. 80% of the number of lower value licks were randomly chosen from each session. For example, if a rat emitted 400 licks for the high concentration sucrose and 200 licks for the low concentration sucrose, then 160 licks would be randomly chosen from each of data type to compare the same number of licks for each lick type. This permutation of taking 80% of the licks was re-sampled 25 times and spectral values were recalculated for each permutation. The maximum ITC value was obtained through calculating the absolute value of ITC values between 2 to 12 Hz within a ∼150 ms window (+1 inter-lick interval) around each lick. The maximum ERSP value was also taken around the same frequency and time window. Then, the average maximum ITC or ERSP value (of the 25x resampled values) for each LFP channel for each rat was saved in a data frame, and each electrode’s maximum ITC and ERSP value for each type of lick (high-value or low-value lick) were used in the ANOVAs for group summaries. Group summary for the peak-to-peak Event-Related Potential (ERP) size recorded the average difference between the maximum and minimum ERP amplitude across all frequencies, using + 1 inter-lick interval window around each lick. The mean ERP size for each electrode for each rat was used in the ANOVAs for group summaries. These analyses were performed for all behavioral variations.

## Acknowledgments

We thank Wambura Fobbs, Alexxai Kravitz, Catherine Stoodley, Steve Wise, and Samantha White for helpful feedback on the manuscript.

---

**Figure 1 - figure supplement 1.**
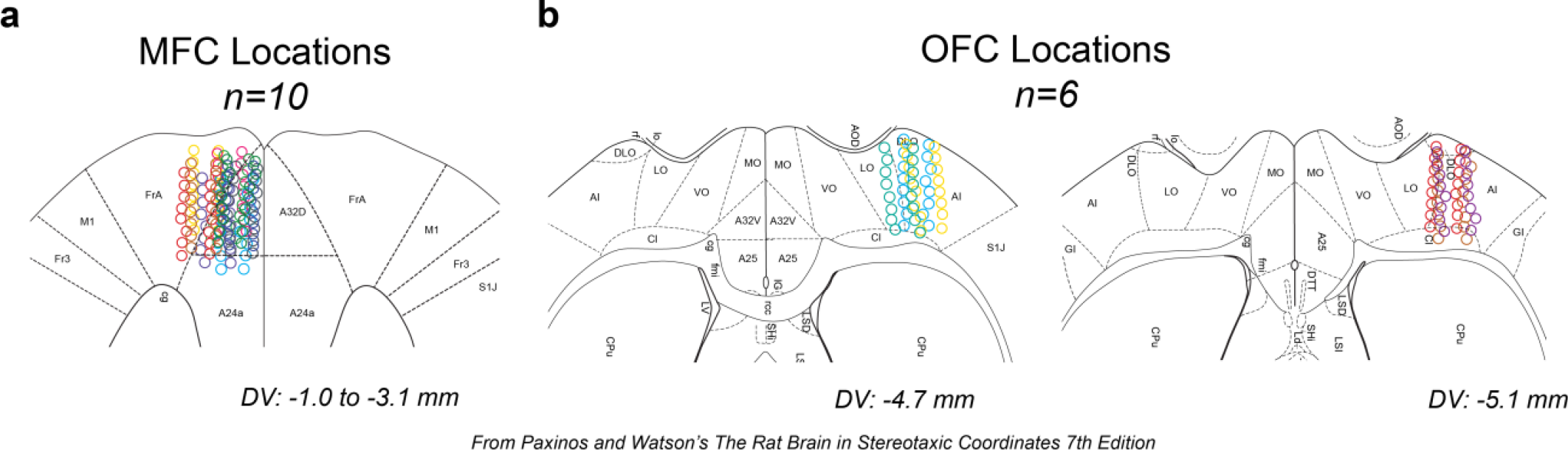
Electrode localization. Locations of all electrodes plotted in the horizontal plane. A. MFC (n=10 rats; 160 electrodes) electrode arrays were localized around area 32 (A32D) and M2 (FrA) from 1 to 3 mm ventral from the brain’s surface. B. OFC (n=6 rats; 96 electrodes) electrode arrays were localized around agranular insular (AI) and lateral orbital (LO) areas of OFC from 4.7 to 5.1 mm ventral from the brain’s surface. Reconstructions were plotted over atlas figures from Paxinos and Watson’s The Rat Brain in Stereotaxic Coordinates, 7^th^ edition (2013).

**Figure 4 - figure supplement 1.**
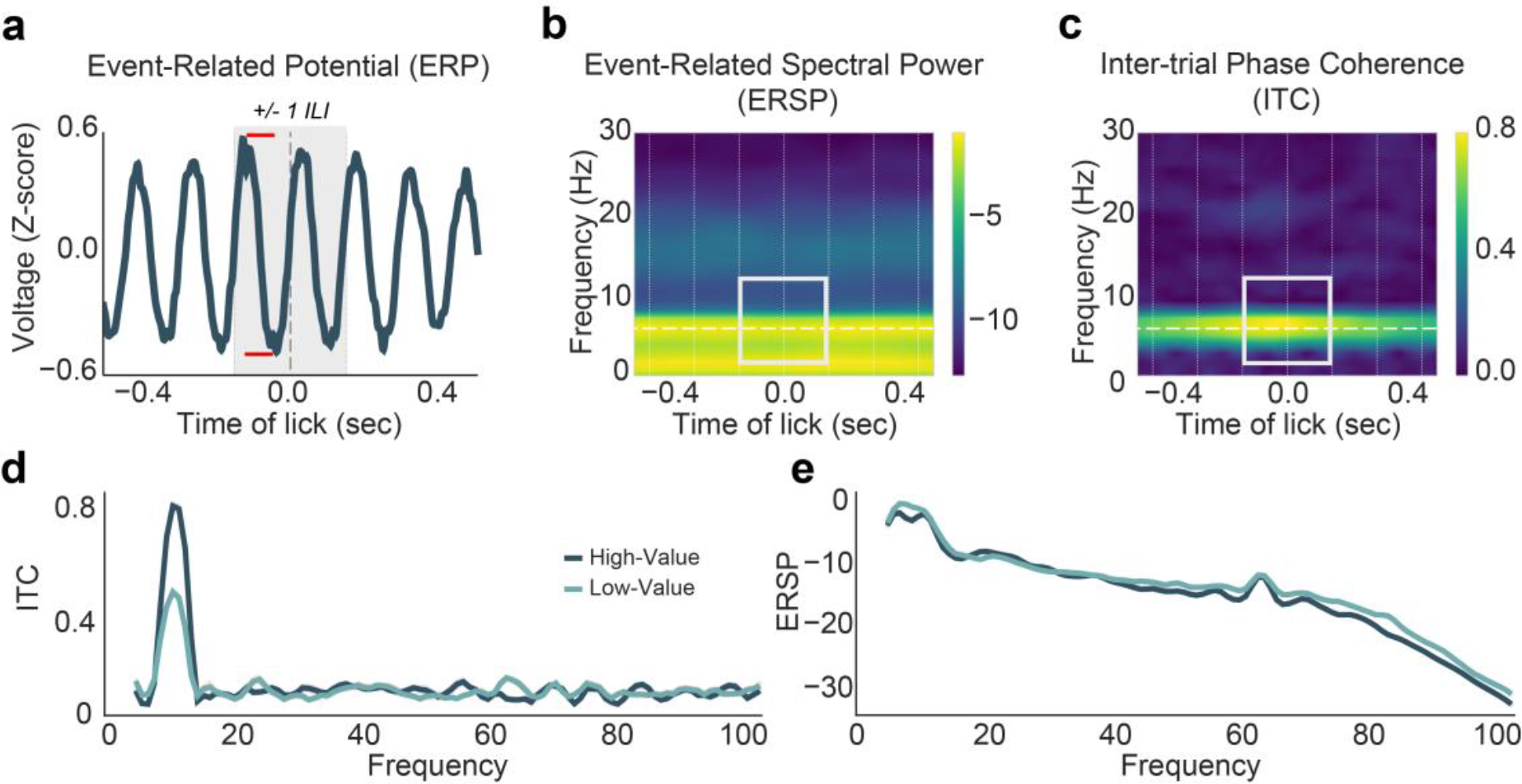
Electrophysiological Measures Used to Assess LFP Activity. A. Event-related potentials (ERPs) were recorded around licks (time 0) after LFP activity was filtered and z-scored. Peak-to-peak analysis was performed on the ERP centered around each lick with a +1 inter-lick interval (ILI) window to calculate the amplitude size (red limits = maximum minus the minimum amplitude of the ERP). B,C. Spectral measures of power (B) and phase (C). Grouped statistics were based on the mean maximum Event-Related Spectral Power (ERSP) and Inter-Trial phase Coherence (ITC) value from 2-12 Hz and around +1 ILI (grey window). Vertical lines denote the rat’s average ILIs. Horizontal line denotes the rat’s median lick rate. D. Maximum ITC values over frequencies from 0-100 Hz from all 16 MFC electrodes from one example rat. E. Maximum ERSP measures over frequencies from 0-100 Hz in all 16 MFC electrodes from one

**Figure 5 - figure supplement 1.**
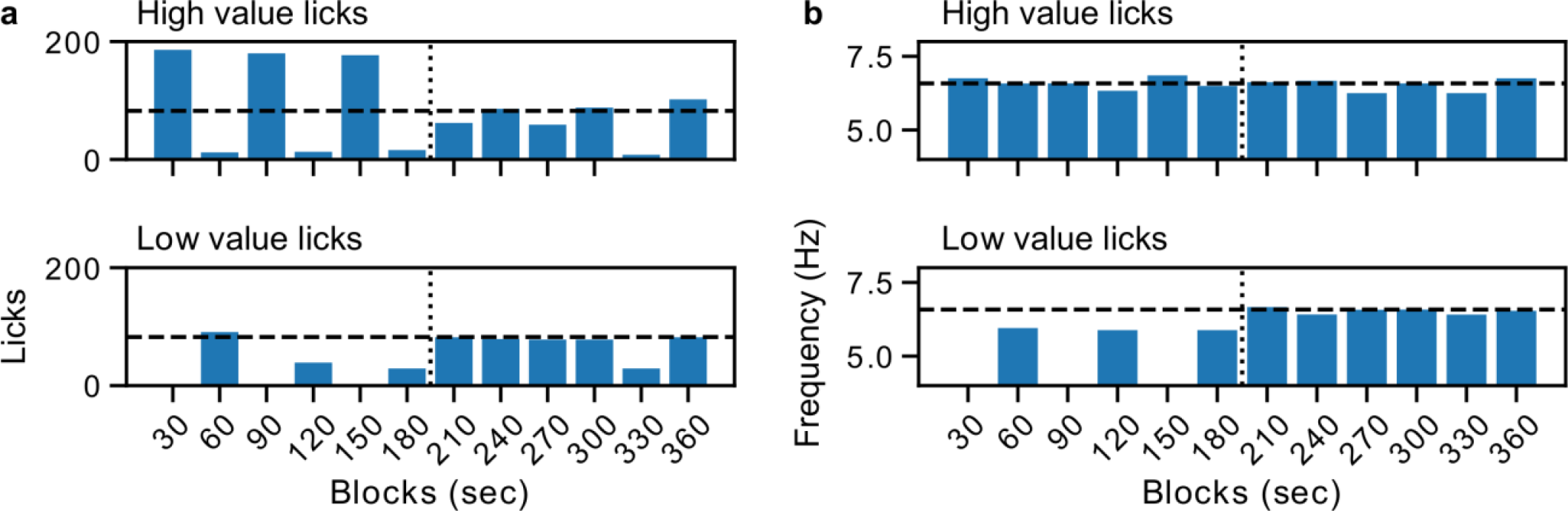
Transitions in licking behavior in the Blocked-Interleaved experiment. Sessions were split into sequential 30 sec windows and the various measures of licking behavior were plotted. An example from one of the rats is shown here. The plots in panel A depict high and low value licks per block, and the dashed line is the mean number of high value licks over all blocks. The plots in panel B show licking frequency for the high and low value fluids, and the dashed line is the median licking rate for the high value fluid over all blocks. There was a clear breakpoint in licks emitted and the licking frequency at the transition from blocked to interleaved presentations of the rewards (vertical dashed line). Licking frequency was lower when rats licked for the lower value fluid when it was presented in the blocked part of the test sessions, and then increased to the same frequency as when they licked for the higher value fluid in the interleaved part of the test sessions. Total licks were higher for the higher value fluid when it was presented in blocks compared to the interleaved part of the session. Licks for the lower value fluid increased starting from the onset of the interleaved part of the session.

**Figure 10 - figure supplement 1.**
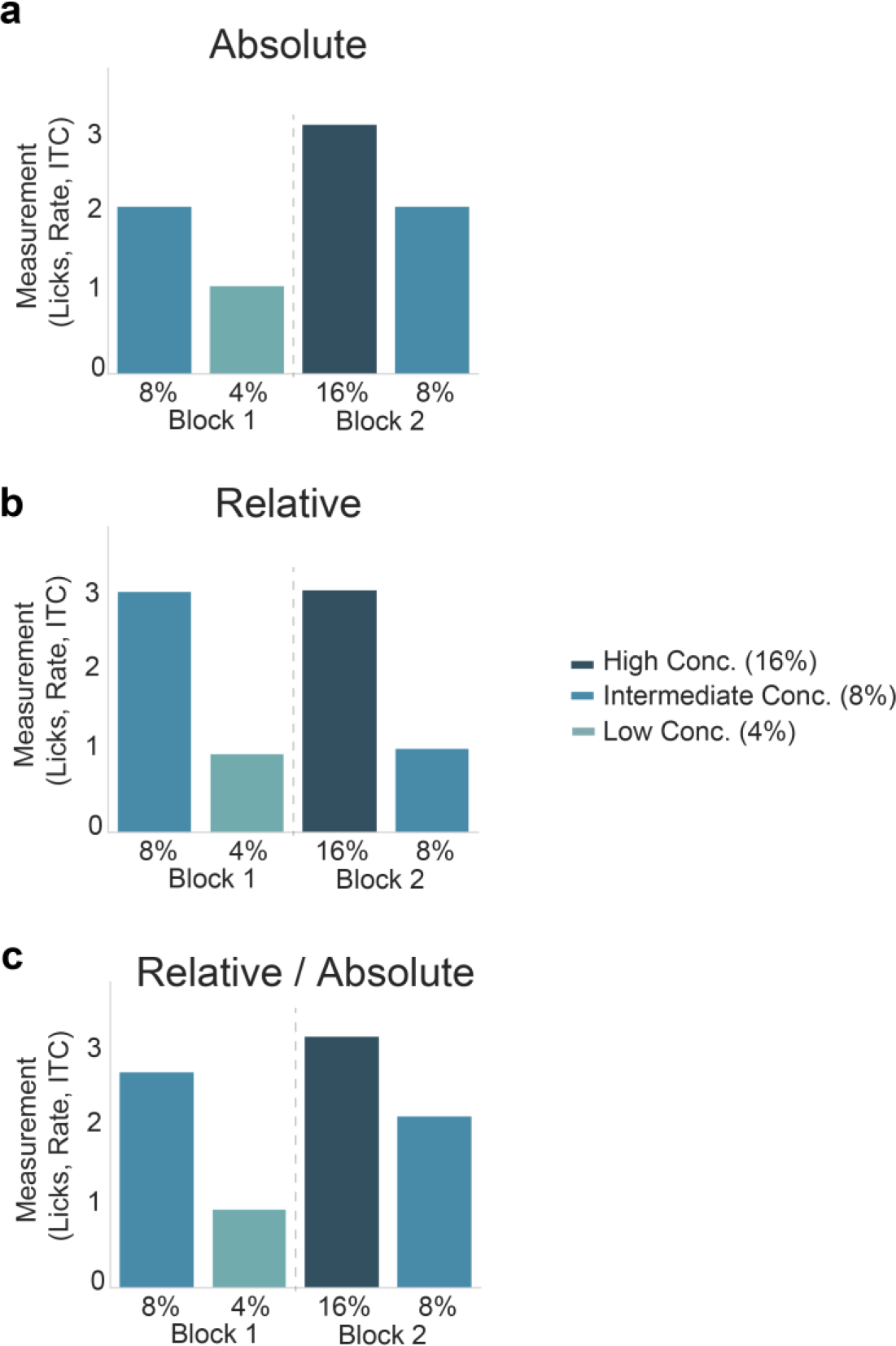
Hypothesis for Relative versus Absolute Encoding of Reward Value. A. If rewards are processed in an absolute manner, we expected to see a graded expression (in lick counts, lick rate, bout duration, or ITC values) of reward value where the high (16%) concentration reward expression is greatest, followed by equal expression of the intermediate (8%) reward and then low expression of the low (4%) concentration reward. B. If rewards are processed in a relative manner, we expected to see a comparative process of rewards, where the “high value” (8% in block 1 or 16% in block 2) are processed similarly, and the focus is on the comparison within each block or context. C. An alternative hypothesis which incorporates a combination of relative and absolute processing of reward value, with partially mixed results of each process in A and B.

**Figure 10 - figure supplement 2.**
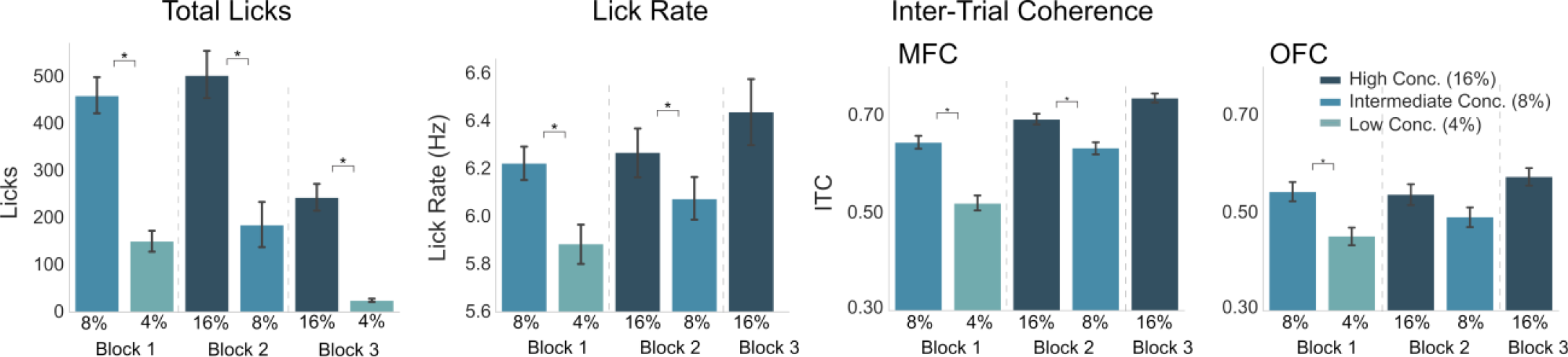
Third Block of Testing in the Three Reward Task. In the third block of the three reward task, rats received access to 16% and 4% sucrose. In this block, licks for 16% sucrose can be compared against licks for both 4% and 8%, and likewise the licks for 8% sucrose can be compared against licks for 4% and 16%. The goal was to attempt to examine if MFC or OFC indeed tracked value in an absolute manner. Overall, rats greatly decreased the number of licks emitted for the 4% sucrose during Block 3 and therefore we could not examine their subsequent electrophysiological findings. However, almost all rats (n=8 in MFC, and n=5 in OFC) emitted at least the minimum criteria of licks for 16% sucrose. Behaviorally, rats emitted more licks for the high value (16%) sucrose in Block 3 as opposed to licks for the low value (4%) sucrose in Block 3 (p<0.005), although there was no difference in total licks emitted for Block 3 high value versus Block 2 intermediate value. There was a significant decrease in the Block 3 high value licks as opposed to Block 2 high value licks (p<0.001). Lick rate was also re-analyzed including Block 3 high value licks, but lick rate for low value (4%) licks could not be analyzed due to a low number of licks not passing criteria. Lick rate for Block 3 high value licks were not significantly different from Block 2 high value licks (p=0.99), nor from Block 2 intermediate licks (p=0.83). Statistics for MFC: Block 3 High value ITCs [95% CI: 0.716, 0.757] were not significantly different from Block 2 High value ITCs (p=0.113) [95% CI: 0.671, 0.715]. Block 3 High value ITCs were significantly increased from both intermediate (8%) value ITCs in Block 1 [95% CI: 0.619, 0.672] and Block 2 [95% CI: 0.610, 0.658] (p<0.001 for both). These findings support the hypothesis of MFC possibly encoding reward value in an absolute manner. Statistics for OFC: Block 3 High value ITCs [95% CI: 0.540, 0.611] were not significantly different from Block 2 High value ITCs (p=0.789) [95% CI: 0.497, 0.580]. Block 3 High value ITCs were significantly increased from Block 2 Intermediate value ITCs (p=0.025) [95% CI for intermediate Block 2: 0.450, 0.535], but Block 3 high value ITCs were not significantly different from Block 1 intermediate ITCs [Block 1 intermediate 95% CI: 0.505, 0.584]. The other comparisons (Block 2 intermediate versus Block 2 high value) were not significant (p=0.339).

## REFERENCES

1. Alexander, W. H., & Brown, J. W. (2011). Medial prefrontal cortex as an action-outcome predictor. Nature Neuroscience, 14(10), 1338–1344. https://doi.org/10.1038/nn.2921

2. Amarante, L. M., Caetano, M. S., & Laubach, M. (2017). Medial frontal theta is entrained to rewarded actions. Journal of Neuroscience, 37(44), 10757–10769. https://doi.org/10.1523/JNeurosci.1965-17.2017

3. Amarante, L. M., Newport, J., Mitchell, M., Wilson, J., & Laubach, M. (2019). An open source syringe pump controller for fluid delivery of multiple volumes. eNeuro, 6(5).https://doi.org/10.1523/eNeuro.0240-19.2019

4. Balleine, B. W., & Dickinson, A. (1998). Goal-directed instrumental action: Contingency and incentive learning and their cortical substrates. Neuropharmacology, 37(4–5), 407–419. https://doi.org/10.1016/S0028-3908(98)00033-1

5. Balleine, B. W., & Dickinson, A. (2000). The effect of lesions of the insular cortex on instrumental conditioning: evidence for a role in incentive memory. Journal of Neuroscience, 20(23), 8954–8964. https://doi.org/10.1523/JNeurosci.20-23-08954.2000

6. Barreiros, I.V.., Panayi, M.C., and Walton, M.E. (2020) Organisation of afferents along the anterior-posterior and medial-lateral axes of the rat OFC. bioRxiv. https://doi.org/10.1101/2020.08.28.272591

7. Bekolay, T., Laubach, M., & Eliasmith, C. (2014). A spiking neural integrator model of the adaptive control of action by the medial prefrontal cortex. Journal of Neuroscience, 34(5), 1892–1902. https://doi.org/10.1523/JNEUROSCI.2421-13.2014

8. Black, R. W. (1968). Shifts in magnitude of reward and contrast effects in instrumental and selective learning: A reinterpretation. Psychological Review, 75(2), 114–126. https://doi.org/10.1037/h0025563

9. Carracedo, L. M., Kjeldsen, H., Cunnington, L., Jenkins, A., Schofield, I., Cunningham, M. O., Davies, C. H., Traub, R. D., & Whittington, M. A. (2013). A neocortical delta rhythm facilitates reciprocal interlaminar interactions via nested theta rhythms. The Journal of neuroscience : the official journal of the Society for Neuroscience, 33(26), 10750–10761. https://doi.org/10.1523/JNEUROSCI.0735-13.2013

10. Cavanagh, J. F., Zambrano-Vazquez, L., & Allen, J. J. B. (2012). Theta lingua franca: A common mid-frontal substrate for action monitoring processes: Omnipresent theta. Psychophysiology, 49(2), 220–238. https://doi.org/10.1111/j.1469-8986.2011.01293.x

11. Cavanagh, J. F., & Frank, M. J. (2014). Frontal theta as a mechanism for cognitive control. Trends in cognitive sciences,18(8), 414–421. https://doi.org/10.1016/j.tics.2014.04.012

12. Collier, G., & Myers, L. (1961). The loci of reinforcement. Journal of Experimental Psychology, 61(1), 57–66. https://doi.org/10.1037/h0048851

13. Collier, G., & Willis, F. N. (1961). Deprivation and reinforcement. Journal of Experimental Psychology, 62(4), 377–384. https://doi.org/10.1037/h0047144

14. Crespi, L. P. (1942). Quantitative variation of incentive and performance in the white rat. The American Journal of Psychology, 55(4), 467. https://doi.org/10.2307/1417120

15. Delorme, A., & Makeig, S. (2004). EEGLAB: An open source toolbox for analysis of single-trial EEG dynamics including independent component analysis. Journal of Neuroscience Methods, 134(1), 9–21. https://doi.org/10.1016/j.jneumeth.2003.10.009

16. Flaherty, C. F. (1982). Incentive contrast: A review of behavioral changes following shifts in reward. Animal Learning and Behavior, 10(4), 409–440. https://doi.org/10.3758/BF03212282

17. Flaherty, C. F., & Rowan, G. A. (1986). Successive, simultaneous, and anticipatory contrast in the consumption of saccharin solutions. Journal of experimental psychology. Animal behavior processes, 12(4), 381–393.

18. Gabbott, P. L. A., Warner, T. A., Jays, P. R. L., & Bacon, S. J. (2003). Areal and synaptic interconnectivity of prelimbic (area 32), infralimbic (area 25) and insular cortices in the rat. Brain Research, 993(1–2), 59–71. https://doi.org/10.1016/j.brainres.2003.08.056

19. Gabbott, P. L. A., Warner, T. A., Jays, P. R. L., Salway, P., & Busby, S. J. (2005). Prefrontal cortex in the rat: Projections to subcortical autonomic, motor, and limbic centers. Journal of Comparative Neurology, 492(2), 145–177. https://doi.org/10.1002/cne.20738

20. Gallagher, M., McMahan, R. W., & Schoenbaum, G. (1999). Orbitofrontal cortex and representation of incentive value in associative learning. Journal of Neuroscience, 19(15), 6610–6614. https://doi.org/10.1523/JNeurosci.19-15-06610.1999

21. Gutierrez, R., Carmena, J. M., Nicolelis, M. A. L., & Simon, S. A. (2006). Orbitofrontal ensemble activity monitors licking and distinguishes among natural rewards. Journal of Neurophysiology, 95(1), 119–133. https://doi.org/10.1152/jn.00467.2005

22. Gutierrez, R., & Simon, S. (2013). Increases in spike timing precision improves gustatory discrimination upon learning. In P. DiLorenzo & J. Victor, Spike Timing (Vol. 20135633, pp. 321–342). CRC Press. https://doi.org/10.1201/b14859-16

23. Horst, N. K., & Laubach, M. (2013). Reward-related activity in the medial prefrontal cortex is driven by consumption. Frontiers in Neuroscience, 7. https://doi.org/10.3389/fnins.2013.00056

24. Hull, C.L. (1943). Principles of Behavior (Vol. 422). New York: Appleton-Century-Crofts.

25. Hulse, S. H., Snyder, H. L., & Bacon, W. E. (1960). Instrumental licking behavior as a function of schedule, volume, and concentration of a saccharine reinforcer. Journal of Experimental Psychology, 60(6), 359–364. https://doi.org/10.1037/h0045841

26. Izquierdo, A. (2017). Functional heterogeneity within rat orbitofrontal cortex in reward learning and decision making. Journal of Neuroscience, 37(44), 10529–10540. https://doi.org/10.1523/JNeurosci.1678-17.2017

27. Jarovi, J., Volle, J., Yu, X., Guan, L., & Takehara-Nishiuchi, K. (2018). Prefrontal theta oscillations promote selective encoding of behaviorally relevant events. eNeuro, 5(6). https://doi.org/10.1523/Eneuro.0407-18.2018

28. Kaplan, J. M., Baird, J. P., & Grill, H. J. (2001). Dissociation of licking and volume intake controls in rats ingesting glucose and maltodextrin. Behavioral neuroscience, 115(1), 188.

29. Kesner, R. P., & Gilbert, P. E. (2007). The role of the agranular insular cortex in anticipation of reward contrast. Neurobiology of Learning and Memory, 88(1), 82–86. https://doi.org/10.1016/j.nlm.2007.02.002

30. Klimesch, W., Sauseng, P., Hanslmayr, S., Gruber, W., & Freunberger, R. (2007). Event-related phase reorganization may explain evoked neural dynamics. Neuroscience and biobehavioral reviews, 31(7), 1003–1016. https://doi.org/10.1016/j.neubiorev.2007.03.005

31. Kopell, N., Kramer, M. A., Malerba, P., & Whittington, M. A. (2010). Are different rhythms good for different functions?. Frontiers in human neuroscience, 4, 187. https://doi.org/10.3389/fnhum.2010.00187

32. Laubach, M., Amarante, L. M., Swanson, K., & White, S. R. (2018). What, if anything, is rodent prefrontal cortex? Eneuro, 5(5). https://doi.org/10.1523/Eneuro.0315-18.2018

33. Laubach, M., Caetano, M. S., & Narayanan, N. S. (2015). Mistakes were made: Neural mechanisms for the adaptive control of action initiation by the medial prefrontal cortex. Journal of Physiology-Paris, 109(1–3), 104–117. https://doi.org/10.1016/j.jphysparis.2014.12.001

34. Levy, D. J., & Glimcher, P. W. (2011). Comparing apples and oranges: using reward-specific and reward-general subjective value representation in the brain. Journal of Neuroscience, 31(41), 14693–14707. https://doi.org/10.1523/JNeurosci.2218-11.2011

35. Levy, D. J., & Glimcher, P. W. (2012). The root of all value: A neural common currency for choice. Current Opinion in Neurobiology, 22(6), 1027–1038. https://doi.org/10.1016/j.conb.2012.06.001

36. Lin, J.-Y., Roman, C., & Reilly, S. (2009). Insular cortex and consummatory successive negative contrast in the rat. Behavioral Neuroscience, 123(4), 810–814. https://doi.org/10.1037/a0016460

37. Lockmann, A., & Tort, A. (2018). Nasal respiration entrains delta-frequency oscillations in the prefrontal cortex and hippocampus of rodents. Brain structure & function, 223(1), 1–3. https://doi.org/10.1007/s00429-017-1573-1

38. Montague, P. R., & Berns, G. S. (2002). Neural economics and the biological substrates of valuation. Neuron, 36(2), 265–284. https://doi.org/10.1016/S0896-6273(02)00974-1

39. Narayanan, N. S., Cavanagh, J. F., Frank, M. J., & Laubach, M. (2013). Common medial frontal mechanisms of adaptive control in humans and rodents. Nature Neuroscience, 16(12), 1888–1895. https://doi.org/10.1038/nn.3549

40. Öngür, D., & Price, J. L. (2000). The organization of networks within the orbital and medial prefrontal cortex of rats, monkeys and humans. Cerebral cortex, 10(3), 206–219. https://doi.org/10.1093/cercor/10.3.206

41. Padoa-Schioppa, C. (2011). Neurobiology of economic choice: A good-based model. Annual Review of Neuroscience, 34(1), 333–359. https://doi.org/10.1146/annurev-neuro-061010-113648

42. Padoa-Schioppa, C., & Assad, J. A. (2006). Neurons in the orbitofrontal cortex encode economic value. Nature, 441(7090), 223–226. https://doi.org/10.1038/nature04676

43. Padoa-Schioppa, C., & Assad, J. A. (2008). The representation of economic value in the orbitofrontal cortex is invariant for changes of menu. Nature Neuroscience, 11(1), 95– 102. https://doi.org/10.1038/nn2020

44. Parent, M. A., Amarante, L. M., Liu, B., Weikum, D., & Laubach, M. (2015a). The medial prefrontal cortex is crucial for the maintenance of persistent licking and the expression of incentive contrast. Frontiers in Integrative Neuroscience, 9. https://doi.org/10.3389/fnint.2015.00023

45. Parent, M. A., Amarante, L. M., Swanson, K., & Laubach, M. (2015b). Cholinergic and ghrelinergic receptors and KCNQ channels in the medial PFC regulate the expression of palatability. Frontiers in behavioral neuroscience, 9, 284. https://doi.org/10.3389/fnbeh.2015.00284

46. Passingham, R. E., & Wise, S. P. (2012). The neurobiology of the prefrontal cortex: Anatomy, evolution, and the origin of insight. Oxford University Press.

47. Paxinos, G., & Watson, C. (2013). The rat brain in stereotaxic coordinates. Seventh Edition. Elsevier Academic Press.

48. Petykó, Z., Gálosi, R., Tóth, A., Máté, K., Szabó, I., Szabó, I., Karádi, Z., & Lénárd, L. (2015). Responses of rat medial prefrontal cortical neurons to pavlovian conditioned stimuli and to delivery of appetitive reward. Behavioural Brain Research, 287, 109–119. https://doi.org/10.1016/j.bbr.2015.03.034

49. Petykó, Z., Tóth, A., Szabó, I., Gálosi, R., & Lénárd, L. (2009). Neuronal activity in rat medial prefrontal cortex during sucrose solution intake: NeuroReport, 20(14), 1235–1239. https://doi.org/10.1097/WNR.0b013e32832fbf30

50. Pratt, W. E., & Mizumori, S. J. Y. (2001). Neurons in rat medial prefrontal cortex show anticipatory rate changes to predictable differential rewards in a spatial memory task. Behavioural Brain Research, 123(2), 165–183. https://doi.org/10.1016/S0166-4328(01)00204-2

51. Rangel, A., Camerer, C., & Montague, P. R. (2008). A framework for studying the neurobiology of value-based decision making. Nature Reviews Neuroscience, 9(7), 545–556.

52. Rangel, A. (2013). Regulation of dietary choice by the decision-making circuitry. Nature Neuroscience, 16(12), 1717–1724. https://doi.org/10.1038/nn.3561

53. Riceberg, J. S., & Shapiro, M. L. (2017). Orbitofrontal cortex signals expected outcomes with predictive codes when stable contingencies promote the integration of reward history. Journal of Neuroscience, 37(8), 2010–2021. https://doi.org/10.1523/JNeurosci.2951-16.2016

54. Saez, R. A., Saez, A., Paton, J. J., Lau, B., & Salzman, C. D. (2017). Distinct roles for the amygdala and orbitofrontal cortex in representing the relative amount of expected reward. Neuron, 95(1), 70–77.e3. https://doi.org/10.1016/j.neuron.2017.06.012

55. Sauseng, P., Klimesch, W., Gruber, W. R., Hanslmayr, S., Freunberger, R., & Doppelmayr, M. (2007). Are event-related potential components generated by phase resetting of brain oscillations? A critical discussion. Neuroscience, 146(4), 1435–1444. https://doi.org/10.1016/j.neuroscience.2007.03.014

56. Schoenbaum, G., & Roesch, M. (2005). Orbitofrontal cortex, associative learning, and expectancies. Neuron, 47(5), 633–636. https://doi.org/10.1016/j.neuron.2005.07.018

57. Schoenbaum, G., Roesch, M. R., Stalnaker, T. A., & Takahashi, Y. K. (2009). A new perspective on the role of the orbitofrontal cortex in adaptive behaviour. Nature Reviews Neuroscience, 10(12), 885–892. https://doi.org/10.1038/nrn2753

58. Sclafani, A., Hertwig, H., Vigorito, M., & Feigin, M. B. (1987). Sex differences in polysaccharide and sugar preferences in rats. Neuroscience and biobehavioral reviews, 11(2), 241–251. https://doi.org/10.1016/s0149-7634(87)80032-5

59. Simon, N. W., Wood, J., & Moghaddam, B. (2015). Action-outcome relationships are represented differently by medial prefrontal and orbitofrontal cortex neurons during action execution. Journal of Neurophysiology, 114(6), 3374–3385. https://doi.org/10.1152/jn.00884.2015

60. Siniscalchi, M. J., Wang, H., & Kwan, A. C. (2019). Enhanced population coding for rewarded choices in the medial frontal cortex of the mouse. Cerebral Cortex, 29(10), 4090–4106.

61. Spence, K.W. (1956). Behavior Theory and Conditioning.

62. Strait, C. E., Blanchard, T. C., & Hayden, B. Y. (2014). Reward value comparison via mutual inhibition in ventromedial prefrontal cortex. Neuron, 82(6), 1357–1366. https://doi.org/10.1016/j.neuron.2014.04.032

63. Sugrue, L. P., Corrado, G. S., & Newsome, W. T. (2005). Choosing the greater of two goods: Neural currencies for valuation and decision making. Nature Reviews Neuroscience, 6(5), 363–375. https://doi.org/10.1038/nrn1666

64. Sul, J. H., Jo, S., Lee, D., & Jung, M. W. (2011). Role of rodent secondary motor cortex in value-based action selection. Nature Neuroscience, 14(9), 1202–1208. https://doi.org/10.1038/nn.2881

65. Swanson, K., Goldbach, H. C., & Laubach, M. (2019). The rat medial frontal cortex controls pace, but not breakpoint, in a progressive ratio licking task. Behavioral neuroscience, 133(4), 385–397. https://doi.org/10.1037/bne0000322

66. Travers, J. B., Dinardo, L. A., & Karimnamazi, H. (1997). Motor and premotor mechanisms of licking. Neuroscience and Biobehavioral Reviews, 21(5), 631–647. https://doi.org/10.1016/S0149-7634(96)00045-0

67. Tremblay, L., & Schultz, W. (1999). Relative reward preference in primate orbitofrontal cortex. Nature, 398(6729), 704–708. https://doi.org/10.1038/19525

68. Uchida, N., & Mainen, Z. F. (2003). Speed and accuracy of olfactory discrimination in the rat. Nature Neuroscience, 6(11), 1224–1229. https://doi.org/10.1038/nn1142

69. van Duuren, E., van der Plasse, G., Lankelma, J., Joosten, R. N. J. M. A., Feenstra, M. G. P., & Pennartz, C. M. A. (2009). Single-cell and population coding of expected reward probability in the orbitofrontal cortex of the rat.. Journal of Neuroscience, 29(28), 8965– 8976. https://doi.org/10.1523/JNeurosci.0005-09.2009

70. van Wingerden, M., Vinck, M., Lankelma, J., & Pennartz, C. M. A. (2010). Theta-band phase locking of orbitofrontal neurons during reward expectancy. Journal of Neuroscience, 30(20), 7078–7087. https://doi.org/10.1523/JNeurosci.3860-09.2010

71. Webber, E. S., Chambers, N. E., Kostek, J. A., Mankin, D. E., & Cromwell, H. C. (2015). Relative reward effects on operant behavior: Incentive contrast, induction and variety effects. Behavioural Processes, 116, 87–99. https://doi.org/10.1016/j.beproc.2015.05.003

72. Weijnen, J. (1998). Licking behavior in the rat: Measurement and situational control of licking frequency. Neuroscience and Biobehavioral Reviews, 22(6), 751–760. https://doi.org/10.1016/S0149-7634(98)00003-7

73. Widmann, A., & Schröger, E. (2012). Filter effects and filter artifacts in the analysis of electrophysiological data. Frontiers in Psychology, 3. https://doi.org/10.3389/fpsyg.2012.00233

74. Womelsdorf, T., Vinck, M., Leung, L. S., & Everling, S. (2010). Selective theta-synchronization of choice-relevant information subserves goal-directed behavior. Frontiers in Human Neuroscience, 4. https://doi.org/10.3389/fnhum.2010.00210

75. Yoshida, A., Taki, I., Chang, Z., Iida, C., Haque, T., Tomita, A., Seki, S., Yamamoto, S., Masuda, Y., Moritani, M., & Shigenaga, Y. (2009). Corticofugal projections to trigeminal motoneurons innervating antagonistic jaw muscles in rats as demonstrated by anterograde and retrograde tract tracing. The Journal of comparative neurology, 514(4), 368–386. https://doi.org/10.1002/cne.22013

